# An RNase Toxin Hijacks Elongation Factor-Tu to Cleave the Ribosomal Sarcin-Ricin Loop

**DOI:** 10.64898/2026.09.18.752725

**Authors:** Julia T. Hespanhol, Taylor M. Blackburn, Kristi Pham, Akanksha Varshney, Gianlucca G. Nicastro, Robson F. de Souza, L. Aravind, Arlen Johnson, Kenneth C. Keiler, Christine M. Dunham, Ethel Bayer-Santos

**Affiliations:** Department of Molecular Biosciences, College of Natural Sciences, University of Texas at Austin, Austin, USA, 78749; Department of Chemistry, Emory University, Atlanta, Georgia, USA, 30322; Division of Intramural Research, National Library of Medicine, National Institutes of Health, Bethesda, Maryland, 20894, USA; Departamento de Microbiologia, Instituto de Ciências Biomédicas, Universidade de São Paulo, São Paulo, Brazil, 05508-900; LaMontagne Center for Infectious Disease, The University of Texas at Austin, Austin, Texas, USA, 78749

**Keywords:** RNase, ribotoxin, sarcin–ricin loop, elongation factor Tu, biological conflict systems

## Abstract

Ribosome-targeting toxins typically recognize and attack their substrates directly. Here, we provide the functional characterization of SRLase1, a widespread member of a recently identified RNase superfamily, and show that it cleaves the sarcin-ricin loop (SRL) of 23S rRNA. Unlike previously described ribosome-targeting toxins, SRLase1 requires the host elongation factor Tu (EF-Tu) for activity. EF-Tu normally engages the ribosome as part of the aminoacyl-tRNA delivery cycle, but SRLase1 instead exploits this factor through a mechanism incompatible with canonical ternary-complex formation. This host-factor-assisted mechanism reveals a distinct strategy for translation inhibition, in which a toxin repurposes a conserved component of the elongation machinery to access a vulnerable ribosomal target. Homologs of SRLase1 are widespread across bacterial phyla and are frequently encoded in conflict-associated genomic loci. Together, these findings define a widespread family of SRL-targeting toxins, expand the mechanistic repertoire of ribosomal antibacterial toxins, and demonstrate how conserved components of the host translation machinery can be co-opted to promote toxin activity.

**Significance statement:** Bacterial competition systems encode thousands of predicted toxin proteins, many of which belong to uncharacterized families. Here, we provide the first functional characterization of the BECR-like RNase superfamily and identify SRLase1 as the founding member of a widespread family of ribotoxins. SRLase1 inhibits protein synthesis by cleaving the sarcin-ricin loop, an essential functional center of the ribosome. Unexpectedly, this activity requires elongation factor Tu (EF-Tu), a conserved translation factor that the toxin hijacks to access and cleave its target. These findings uncover a new mechanism of ribosome inactivation, expand the functional diversity of RNase toxins, and demonstrate how conserved host proteins can be co-opted to promote toxin activity during microbial conflict.

## Introduction

To gain an advantage over competitors, many bacteria deploy protein toxins that damage essential cellular processes in neighboring cells. These toxins can be delivered through multiple antagonistic systems, including the type VI secretion system, and typically function together with cognate immunity proteins that protect producer cells from self-intoxication (1, 2). Among the most widespread weapons in bacterial warfare are polymorphic toxins, a large group of modular proteins composed of conserved toxic domains linked to diverse trafficking/secretion modules (1). This modular organization promotes extensive diversification of antibacterial activities, generating a vast repertoire of effectors that target essential cellular processes in competing cells (1).

Translation is a central cellular process, making it a frequent target during interbacterial competition. Many antibacterial systems inhibit protein synthesis through RNase toxins that attack distinct components of the translational apparatus (3–5). Some of these toxins have unspecific RNase activity (6–8) whereas others specifically target transfer RNAs (tRNAs) required for amino acid delivery to the ribosome (9–11). Direct attacks on rRNA appear comparatively rare. Although colicin E3 cleaves the 16S rRNA decoding center (12) and VapC20 from toxin-antitoxin (TA) systems targets the sarcin-ricin loop (SRL) of 23S rRNA (13, 14), the large subunit rRNA has not been reported as a target of antibacterial toxins deployed during interbacterial competition.

Among translation-targeting toxins, a large and diverse group of RNases shares a common structural scaffold despite exhibiting little or no detectable sequence similarity. This group, known as the BECR fold, was originally defined from structural similarities between Barnase, EndoU, Colicin D/E5, and RelE toxins (1). Members of this broader BECR superfamily are frequently associated with polymorphic toxin systems and have evolved diverse substrate specificities (15–19). However, many predicted BECR proteins remain functionally uncharacterized, limiting our understanding of how this fold has diversified to attack different steps of the translation process.

Unlike mRNAs and tRNAs, rRNAs are embedded within a large ribonucleoprotein assembly, creating unique challenges for toxin access and substrate recognition (20, 21). Successful and efficient attacks on rRNA therefore require mechanisms capable of identifying specific functional centers within the assembled ribosome. One particularly vulnerable target is the SRL of the 23S rRNA, a highly conserved ribosomal element that is essential for recruiting and activating elongation factor GTPases that drive protein synthesis (22). How antibacterial RNases engage these highly structured substrates remains poorly understood. Defining the molecular strategies used to target this sequence in active ribosomes is therefore essential for understanding how translation-targeting toxins achieve specificity and potency during bacterial warfare.

Recent comparative genomic analyses uncovered a widespread family of predicted RNases with a distinct structural scaffold that is reminiscent of the classical BECR fold, herein named BECR-like (23, 24). However, the activity and biological targets of these proteins remain unknown. Here, we characterize SRLase1, a representative member of this group, and show that it inhibits translation through specific cleavage of the SRL of 23S rRNA. We further demonstrate that SRLase1 requires elongation factor Tu (EF-Tu) for activity on ribosomes, revealing a host-factor-assisted mechanism for ribosome attack. These findings establish SRLase1 as the founding member of a widespread family of ribosome-targeting antibacterial toxins and uncover an EF-Tu-dependent strategy for translation inhibition.

## Results

### SRLase1 defines a broadly distributed polymorphic toxin family

To investigate whether predicted BECR-like proteins function as antibacterial toxins, we selected a representative protein from *Salmonella enterica* serovar Oslo (*S.* Oslo) that was previously annotated as STox4 (23). We refer to this protein as SRLase1 to reflect the SRL cleavage activity demonstrated below. To determine whether SRLase1 represents a *Salmonella*-specific effector or a member of a broader toxin family, we searched for SRLase1 homologs and analyzed their taxonomic distribution and genomic context. SRLase1 homologs were identified across diverse bacterial phyla, including Pseudomonadota, Myxococcota, Actinomycetota, Bacillota, Bacteroidota, Planctomycetota, Verrucomicrobiota, Chloroflexota, Fibrobacterota, and Desulfobacterota, and were also detected in archaeal genomes (Fig. 1A–C; Dataset S1). Many homologs were encoded within modular genomic organizations characteristic of polymorphic toxin systems, in which a conserved toxic domain is fused to variable trafficking modules and is paired with a nearby immunity gene (1). These architectures included fusions with PAAR, LXG, PT-VENN, and other delivery-associated domains (Fig. 1B–C; Fig. S1). Thus, SRLase1 proteins define a broadly distributed family of BECR-like toxin domains embedded in diverse conflict-associated contexts.

**Fig 1.**
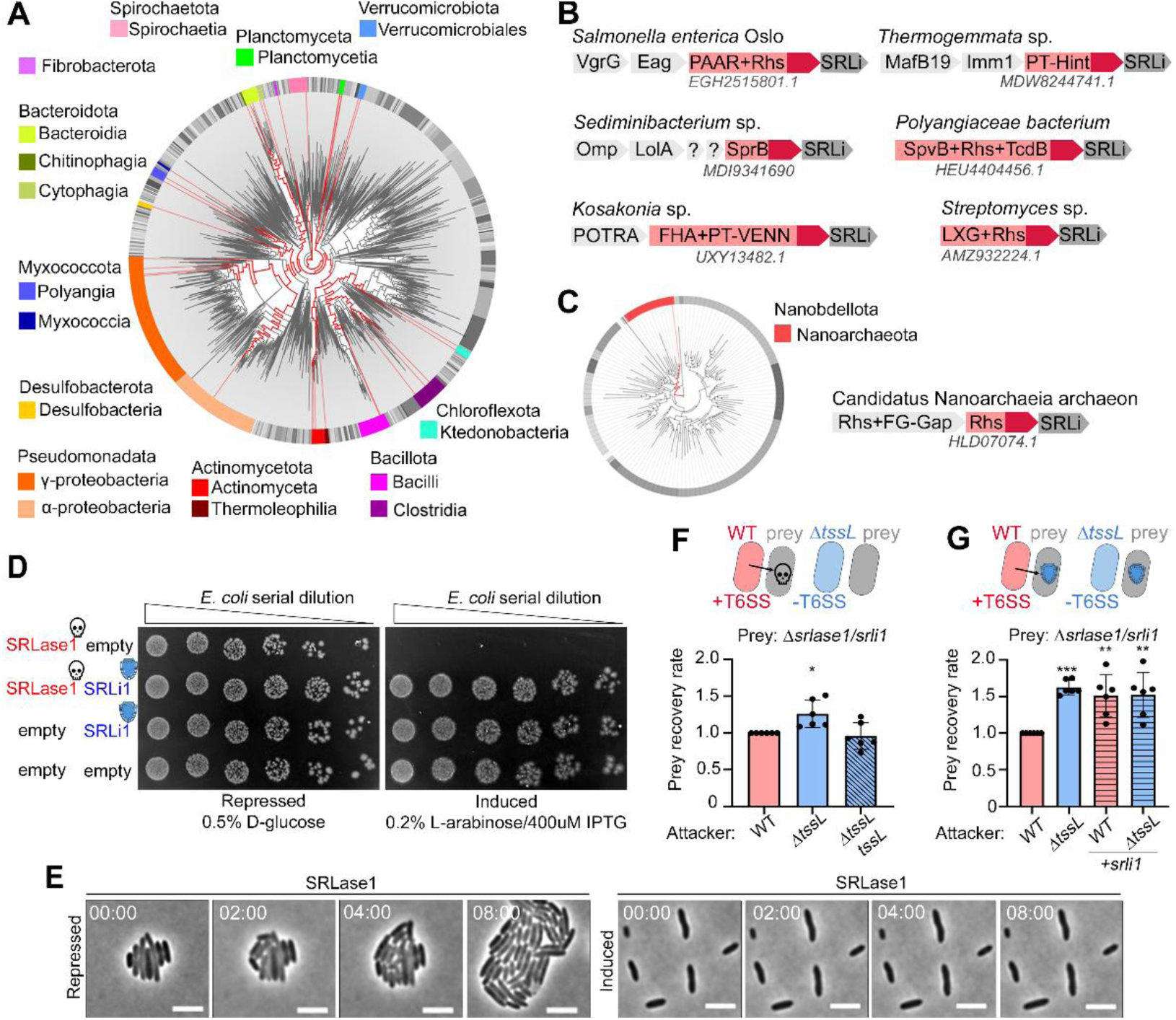
SRLase1 is a widespread polymorphic toxin used in interbacterial competition. **(a)** Bacterial and **(c)** archaeal trees generated using AnnoTree, with nodes marked in red indicating taxa encoding SRLase1 homologues. Inner nodes represent orders, while classes are shown in the outer ring, with major classes color-coded. **(b-c)** Representative genomic neighborhoods of SRLase1 homologs are shown, with SRLase1 homologs in red and SRLi1 in dark gray. The C-terminal SRLase1 domains are indicated by dark red arrow. **(d)** *E. coli* toxicity assay showing serial dilutions of cells carrying pBRA-SRLase1 and pEXT22-SRLi1. Images are representative of three independent experiments. **(e)** Time-lapse microscopy of *E. coli* carrying pBRA-SRLase1 grown under repressing (0.2% D-glucose) or inducing (0.2% L-arabinose) conditions. Scale bar, 5 μm. Time is shown as hh:mm. **(f-g**) Top, schematic of the T6SS-dependent bacterial competition assay. Attacker strains were *S.* Oslo wild-type (WT, red), Δ*tssL* (blue), or Δ*tssL* complemented (blue). T6SS system is represented by the arrow. The prey strain lacked Δ*srlase1-srli1* (gray). **(f)** Competitions between wild-type, Δ*tssL* and Δ*tssL* complemented (blue with diagonal lines) against the prey. **(g)** Competition assay using prey strains lacking *srli1* or complemented with *srli1* (pattern). Data represents the mean ± SD of six independent experiments. Statistical significance was assessed by one-way ANOVA followed by Dunnett’s multiple-comparison test. *p<0.05, and ***p < 0.001. Abbreviations: SpvB, *Salmonella* plasmid virulence toxin B; Eag, effector-associated gene; OmpH, outer membrane protein H; LolA, outer-membrane lipoprotein carrier protein; FG-GAP, phenylalanine–glycine motif glycine–alanine–proline β-propeller domain; PT-Hint, pre-toxin Hint protease domain; POTRA, polypeptide-transport-associated domain; FHA, filamentous haemagglutinin; PT-VENN, pre-toxin VENN.

In *S. Oslo*, SRLase1 is encoded within the type VI secretion system (T6SS) locus and displays a canonical polymorphic toxin organization, with an N-terminal PAAR trafficking domain and a predicted C-terminal BECR-like toxic domain (Fig. 1B). Immediately downstream, a small open reading frame encodes a predicted cognate immunity protein, here termed SRLi1 (Fig. 1D). This organization provided a genetically tractable model to test whether SRLase1 functions as an antibacterial toxin and whether SRLi1 protects against its activity.

To assess toxicity, we expressed the C-terminal toxic domain of SRLase1 immediately downstream of the conserved self-cleavage motif (DPLGL) (25) in *Escherichia coli* under the control of an arabinose-inducible promoter. Induction of SRLase1 resulted in strong growth inhibition, whereas co-expression of SRLi1 fully restored growth (Fig. 1D), demonstrating that SRLase1 and SRLi1 constitute a functional toxin–immunity pair. Despite severe growth inhibition, SRLase1-intoxicated cells maintained normal rod morphology and did not undergo lysis as determined by time-lapse microscopy (Fig. 1E; Movies S1–S2). These observations indicate that SRLase1 does not compromise envelope integrity and are consistent with a mechanism distinct from membrane-disrupting or cell wall-targeting toxins.

To determine whether SRLase1 contributes to interbacterial antagonism, we performed competition assays using *S.* Oslo as a model delivery system (Fig. 1F, 1G). A prey strain lacking the *srlase1/srli1* locus exhibited reduced survival when challenged with wild-type attackers, whereas prey recovery increased when competed against a T6SS-deficient Δ*tssL* mutant (Fig. 1F). Complementation of *tssL* restored prey killing to wild-type levels, confirming that toxin delivery requires a functional T6SS. Expression of SRLi1 in prey cells significantly increased survival during competition (Fig. 1G), directly linking the observed fitness defect to SRLase1 intoxication. Together, these results establish SRLase1 as an antibacterial polymorphic toxin whose activity can be delivered by the *S. enterica* T6SS and is specifically neutralized by its cognate immunity protein.

### SRLase1 cleaves the SRL of 23S rRNA and inhibits translation

SRLase1 was previously assigned to a recently recognized BECR-like group of RNases (23, 24). Canonical BECR proteins are built around a conserved architecture consisting of an N-terminal α-helix followed by a four-stranded sheet in the form of a β-meander, whose exposed face houses the catalytic center (1). BECR-like RNases have a comparable exposed β-sheet, which presents the active site, but is distinguished by the addition of a β-strand at the C-terminal edge of the sheet (Fig. 2A, Fig. S2) (24). In some members, the sheet is further expanded by an additional β-strand N-terminal to the core α-helix, which packs parallel to the last strand of the fold. Across this superfamily, the β-strands topologically equivalent to β1 and β2 of canonical BECRs exhibit varying degrees of degeneration. In contrast, the remainder of the β-sheet is strongly conserved across the BECR-like superfamily, including the strands equivalent to β3 and β4 of canonical BECR domains, the C-terminal β5 strand, and, when present, the additional N-terminal strand (Fig. S2). In addition, sequence analysis further revealed pronounced variability in two surface exposed regions: the segment between the α-helix and β3 and the loop connecting β3 and β4. These regions form Loop1 and Loop2, which are distinctive features of the BECR-like superfamily (Fig. 2A, Fig. S2). Thus, while retaining several features reminiscent of the classical BECR RNases, the BECR-like superfamily possesses a distinct structural core, leaving open whether its similarity to canonical BECR proteins reflects deep divergence from a common ancestor or convergent evolution (24).

**Fig 2.**
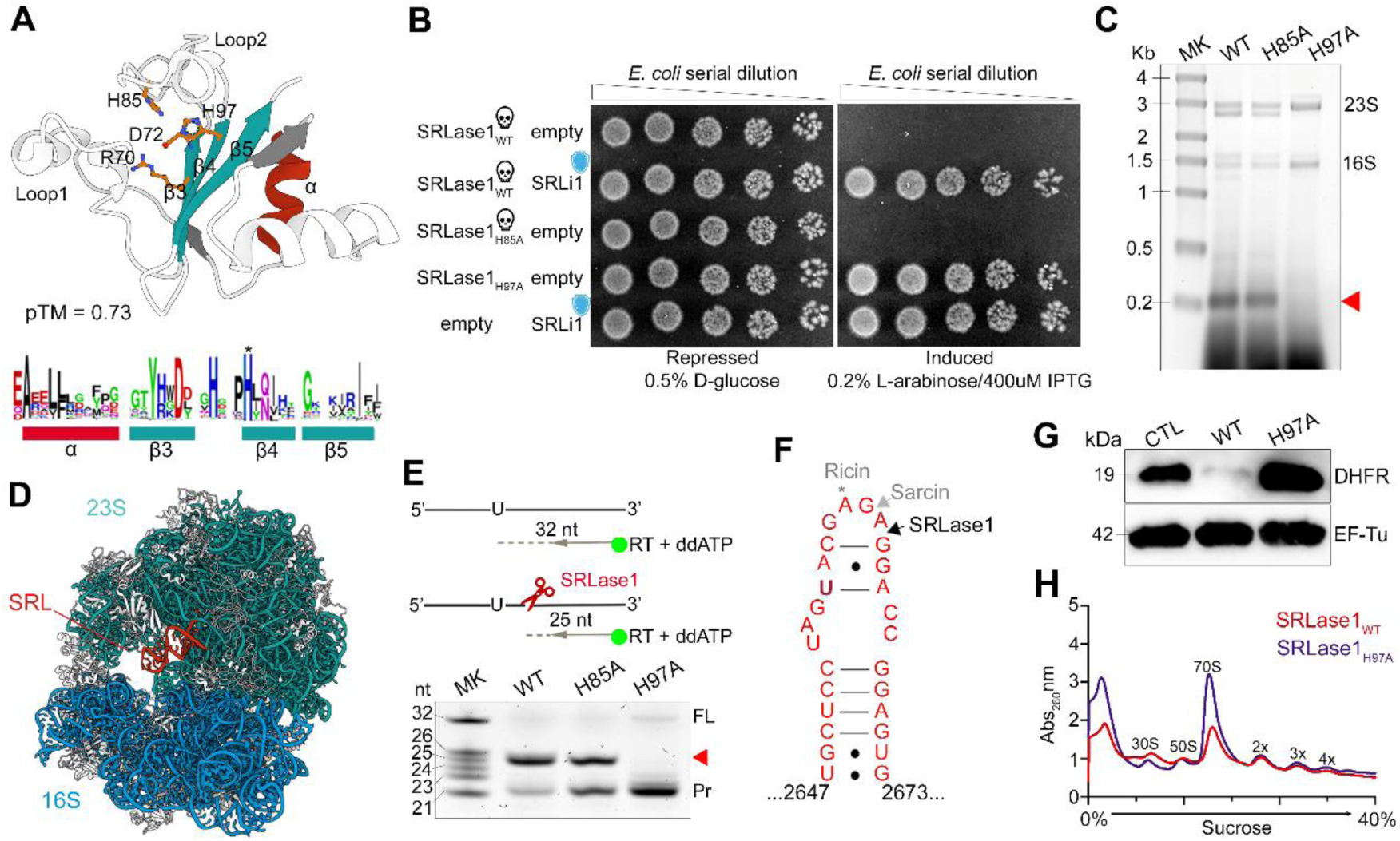
SRLase1 cleaves the SRL of 23S rRNA. **(a)** AlphaFold3 prediction of SRLase1 highlighting a BECR-like architecture. The N-terminal α-helix is shown in red, pre-β strands are in gray, and the central β-sheet in cyan. Conserved residues are shown as orange sticks, and a sequence logo derived from SRLase1 homologs is shown below. **(b)** *E. coli* toxicity assay showing serial dilutions of cells carrying pBRA-SRLase1 constructs and pEXT22-based immunity plasmids. Images are representative of three independent experiments. **(c)** Analysis of total RNA extracted from *E. coli* following expression of the indicated SRLase1 variants. The characteristic ∼200-nt cleavage product generated by SRLase1 is indicated (red arrow). **(d)** Structure of the *E. coli* 70S ribosome (PDB 4V4Q), with 23S rRNA shown in light blue and 16S rRNA shown in dark blue. The sarcin–ricin loop (SRL) is highlighted in red. **(e)** Mapping of the SRLase1 cleavage site by primer extension. Top, schematic of the assay. Bottom, denaturing PAGE of cDNA products generated from RNA isolated from *E. coli* expressing the indicated SRLase1 variants. FAM-labeled primer (21 nt) is labelled as Pr, truncated product generated by SRL cleavage (25 nt) is indicated by red arrowhead, and full-length product (32 nt) by FL. **(f)** Secondary structure of the SRL. The SRLase1 cleavage site between A2662 and G2663 is indicated by a black arrow. The canonical α-sarcin cleavage site between G2661 and A2662 is shown by a gray arrow, and the ricin depurination site (A2660) is indicated by a gray asterisk. **(g)** *In vitro* translation assay. Translation of a DHFR–FLAG reporter was monitored by western blot. Images are representative of three independent experiments. **(h)** Ribosome profiles of *E. coli* expressing SRLase1_WT_ or SRLase1_H97A_. Absorbance at 260 nm is plotted across sucrose gradients. Positions of ribosomal subunits and 70S monosomes are indicated.

To identify residues that might support catalysis within this distinct structural framework, we examined conservation across SRLase1 homologs. BECR-fold RNases generally catalyze substrate cleavage through acid-base chemistry mediated by a small set of conserved charged residues positioned around the active site (15, 17, 26). Histidine residues frequently function as general acids or bases, whereas acidic residues contribute to substrate positioning and stabilization of catalytic intermediates (1, 15, 27, 28). A multiple sequence alignment of SRLase1 homologs revealed four highly conserved residues clustered around the predicted catalytic center: Arg70, Asp72, His85, and His97 (Fig. 2A). To test their functional importance, we generated alanine substitutions for the two histidines. Toxicity assays showed that substitution of H97A completely abolished growth inhibition (Fig. 2B). However, under conditions of reduced repression (0.2% glucose), H85A exhibited a modest but reproducible reduction in toxicity relative to the wild-type protein (Fig. S3), which was still toxic, suggesting that His85 contributes to efficient toxin activity but is not strictly required. Both proteins accumulated to similar levels upon induction (Fig. S4), indicating that loss of activity was not due to impaired protein expression. These results identify His97 as a critical residue for SRLase1 function, while suggesting a more subtle contribution of His85.

To determine whether SRLase1 functions as an RNase, we analyzed total RNA isolated from *E. coli* following toxin induction. Expression of wild-type SRLase1 produced a striking and highly reproducible RNA cleavage pattern characterized by the appearance of a discrete ∼200-nucleotide fragment together with additional species migrating near the 23S rRNA band (Fig. 2C). These products were also observed following expression of the partially toxic SRLase1_H85A_ variant but were absent in cells expressing the inactive SRLase1_H97A_ mutant, demonstrating that their formation depends on SRLase1 catalytic activity. In contrast, neither plasmid degradation nor induction of the SOS response was detected (Fig. S5), suggesting that SRLase1 acts as a dedicated ribonuclease.

The size of the predominant cleavage fragment was unexpected. Its migration pattern closely resembled the characteristic α-fragment generated by cleavage of the SRL, a highly conserved structure located within Helix 95 of the 23S rRNA (Fig. 2D). The SRL forms the principal interaction site for translational GTPases, including EF-Tu (29), EF-G (30) and RF3 (31), and is essential for efficient protein synthesis. This vulnerability is exploited by classical ribotoxins such as α-sarcin, a toxin produced by the filamentous fungus *Aspergillus giganteus*, which cleaves the phosphodiester bond between G2661 and A2662; and ricin, a plant toxin from *Ricinus communis*, which depurinates A2660 within the same loop (32–34). The similarity between the SRLase1 cleavage pattern and the characteristic α-fragment prompted us to test whether SRLase1 targets this ribosomal element.

To identify the cleavage site, we performed a modified primer extension assay using RNA isolated from intoxicated *E. coli* cells. A fluorophore-labeled primer annealing downstream of the SRL was used for cDNA synthesis with ddATP and generates a 32-nt cDNA product when the 23S rRNA template remains intact, whereas cleavage within the SRL produces a shorter product (Fig. 2E). RNA from cells expressing SRLase1_WT_ or SRLase1_H85A_ generated a truncated 25-nt product, whereas RNA from cells expressing the inactive SRLase1_H97A_ mutant yielded only the full-length species (Fig. 2E). The size of the truncated product mapped the cleavage event to the phosphodiester bond between A2662 and G2663 of the 23S rRNA. This site is immediately adjacent to, but distinct from, that targeted by α-sarcin (Fig. 2F). These findings demonstrate that SRLase1 specifically cleaves the SRL of 23S rRNA in intoxicated cells and provides the basis for renaming this toxin SRLase1.

To gain insight into the possible catalytic mechanism of this BECR-like RNase, we examined AlphaFold3 models (35) of SRLase1 bound to the SRL (and EF-Tu·GDP, see below). In the predicted complex, Asp72 is positioned adjacent to the 2′-OH of SRL A2662, supporting a role as the catalytic general base, while His85, His97, and Asn99 flank Asp72 and may contribute to positioning or tuning the catalytic environment of the active site (Fig. S6). Arg70 is located near the attacking 2′-OH and could facilitate catalysis stabilizing the negatively charged transition state (36). Together, the AF3 model and experimental data support a metal-independent RNase mechanism in which Asp72 activates the 2′-OH for inline attack on the phosphodiester bond between A2662 and G2663, generating 2′,3′-cyclic phosphate and 5′-OH products.

Consistent with disruption of SRL function, addition of purified recombinant SRLase1 to a reconstituted *in vitro E. coli* translation system strongly inhibited protein synthesis, whereas reactions supplemented with buffer alone or the catalytic mutant SRLase1_H97A_ remained active (Fig. 2G). RNA recovered from translation reactions containing SRLase1 exhibited the characteristic SRL cleavage product (Fig. S7), directly linking rRNA cleavage to translational arrest. Consistent with these observations, sucrose-gradient analysis of ribosomes isolated from intoxicated *E. coli* cells revealed a marked reduction in the 70S monosome population relative to cells expressing SRLase1_H97A_, indicating widespread disruption of functional ribosomes (Fig. 2H). Together, these findings demonstrate that SRL cleavage by SRLase1 directly compromises prokaryotic ribosome function and provides the mechanistic basis for translation inhibition.

### EF-Tu is required for SRLase1 activity on ribosomes

Classical SRL-targeting toxins access and cleave the ribosome directly without requiring accessory host factors. We therefore asked whether SRLase1 employs a similar strategy or instead depends on host factors for activity. During purification of recombinant SRLase1, we consistently observed co-elution of an endogenous *E. coli* protein (Fig. 3A; Fig. S8). Mass spectrometry identified this protein as EF-Tu, an essential translation factor that delivers aminoacyl-tRNAs to the ribosome and contacts the SRL during the elongation cycle (22, 29, 33). Because EF-Tu directly engages the SRL during translation, its association with SRLase1 raised the possibility that the toxin exploits this host factor to recognize, access, and/or efficiently cleave its ribosomal target.

**Fig 3.**
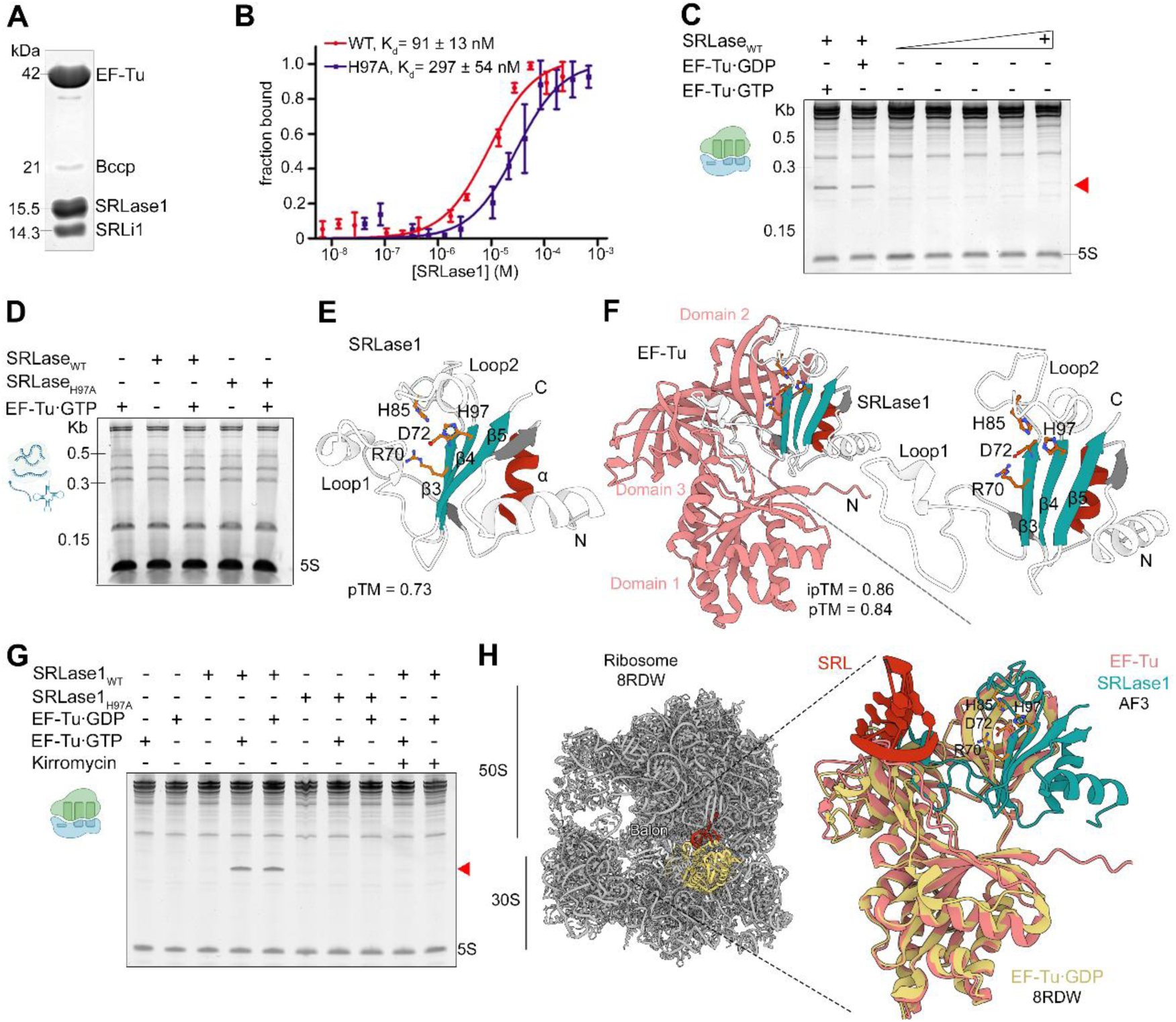
EF-Tu binds SRLase1 and is required for ribosome cleavage. **(a)** Co-elution of recombinant SRLase1 with endogenous EF-Tu during purification of SRLase1–SRLi1 complex. **(b)** Microscale thermophoresis analysis of SRLase1 binding to EF-Tu. Fluorescently labeled EF-Tu was incubated with increasing concentrations of SRLase1_WT_ (red) or SRLase1_H97A_ (blue). Data points represent the mean ± SD of three independent experiments. Dissociation constants were determined by nonlinear regression. **(c)** EF-Tu-dependent cleavage of ribosomal RNA by SRLase1. Purified *E. coli* ribosomes were incubated with SRLase1_WT_ in the presence or absence of EF-Tu·GTP or EF-Tu·GDP, or with increasing concentrations of SRLase1. RNA was extracted and analyzed by denaturing PAGE. The SRL cleavage fragment is indicated (red arrow). **(d)** Total *E. coli* RNA incubated with SRLase1 in the presence of EF-Tu. **(e)** AlphaFold3 prediction of SRLase1 alone. **(f)** AlphaFold3 prediction of SRLase1-EF-Tu complex. **(g)** Ribosome cleavage assay using purified *E. coli* ribosomes incubated with SRLase1_WT_ or SRLase1_H97A_ in the presence or absence of EF-Tu and kirromicyn. Images are representative of three independent experiments. **(h)** Overlay of EF-Tu·GDP bound to ribosome with the AlphaFold3 SRLase1-EF-Tu complex.

To quantify this interaction, we measured SRLase1 binding to EF-Tu by microscale thermophoresis (MST). SRLase1 bound EF-Tu in a concentration-dependent manner, with SRLase1_WT_ exhibiting an apparent dissociation constant of approximately 90 nM (Fig. 3B).

The catalytically inactive SRLase1_H97A_ mutant also bound EF-Tu, but with lower affinity, with an apparent dissociation constant of approximately 300 nM. These results demonstrate that SRLase1 forms a high-affinity complex with EF-Tu and suggest that the catalytic region contributes, directly or indirectly, to optimal EF-Tu engagement.

We next asked whether EF-Tu is required for SRLase1 activity on ribosomes. Purified *E. coli* 70S ribosomes were incubated with SRLase1_WT_ in the absence or presence of EF-Tu loaded with either GTP or GDP. SRL cleavage was detected only when EF-Tu was included in the reaction, and both EF-Tu·GTP and EF-Tu·GDP supported cleavage (Fig. 3C). Increasing the concentration of SRLase1 alone did not induce significant cleavage, indicating that EF-Tu is not simply enhancing an inefficient reaction but is required for toxin activity under these conditions.

To determine whether SRLase1 cleavage is broadly directed toward the conserved SRL sequence or instead exhibits specificity for bacterial ribosomes, we incubated purified SRLase1 and EF-Tu in a eukaryotic *in vitro* translation system. No detectable cleavage product was observed nor was protein synthesis hindered (Fig. S9). Because the SRL is highly conserved across bacteria and eukaryotes, these results indicate that sequence conservation alone is insufficient for substrate recognition and suggest that SRLase1 requires specific features of the bacterial ribosome for activity.

To analyze whether SRLase1 can cleave RNA outside the context of intact ribosomes, we incubated purified SRLase1 with total *E. coli* RNA in the presence or absence of EF-Tu. No RNA degradation was observed under these conditions (Fig. 3D), whereas SRLase1 readily cleaved the SRL when intact ribosomes and EF-Tu were present (Fig. 3C). Thus, SRLase1 does not behave as a nonspecific RNase toward free RNA. Instead, its activity is restricted to the ribosomal context and depends on EF-Tu.

To explore how EF-Tu might promote SRLase1 activity, we used AlphaFold3 to model the SRLase1–EF-Tu complex (Fig. 3E, 3F; Fig. S10). The predicted model placed SRLase1 at the interface between EF-Tu domains II and III, near the region that binds aminoacyl-tRNA (37) and yielded confidence scores consistent with a plausible protein–protein interface (ipTM=0.9; pTM=0.86). The model further suggested that EF-Tu binding induces a conformational rearrangement in SRLase1, including displacement of the variable loops that exposes the catalytic pocket. Together, these structural predictions raise the possibility that EF-Tu promotes SRLase1 activity by remodeling the toxin active site rather than serving solely as a ribosome-targeting factor.

Consistent with the predicted overlap between the SRLase1-binding surface and the aminoacyl-tRNA binding region of EF-Tu, we performed native gel-shift assays to test whether SRLase1 could bind EF-Tu simultaneously with tRNA as part of the EF-Tu·GTP·aa-tRNA ternary complex (Fig. S11). Addition of SRLase1 disrupted ternary-complex formation, and promoted dissociation of tRNA from EF-Tu. These results suggest that SRLase1 engages EF-Tu in a state that is incompatible with stable ternary-complex formation. Because several previously characterized BECR-fold contact-dependent inhibition (CDI) toxins use EF-Tu to facilitate cleavage of tRNA substrates (11, 28, 38), we asked whether SRLase1 similarly targets cellular tRNAs. Analysis of total RNA by PAGE did not reveal detectable tRNA degradation following SRLase1 intoxication (Fig. S12). These findings indicate that SRLase1 does not function as a tRNase and support the conclusion that the SRL is its primary physiological target.

The AlphaFold3 model predicted that SRLase1 binds EF-Tu in a conformation most similar to the open GDP-bound state (Fig. 3F). However, both EF-Tu·GDP and EF-Tu·GTP supported SRL cleavage in our reconstituted assay (Fig. 3C), indicating that toxin activity is not restricted to a single nucleotide-bound form. We therefore asked whether the conformational flexibility of EF-Tu, rather than its nucleotide state, is required for SRLase1 activity. Ribosome cleavage reactions were performed in the presence or absence of kirromycin, an antibiotic that locks EF-Tu in a GTP-like closed conformation by binding between domains I and III (39). Kirromycin abolished SRL cleavage despite the presence of EF-Tu (Fig. 3G), indicating that SRLase1 activity requires EF-Tu conformational flexibility. Together with the AlphaFold3 model, these results suggest that the GDP-bound open conformation of EF-Tu is the most relevant state for SRLase1 activity. While cleavage can occur in the presence of either GTP- or GDP-loaded EF-Tu, the requirement for conformational flexibility indicates that productive toxin activity likely depends on the adoption of an open EF-Tu conformation that promotes formation of a catalytically competent SRLase1-ribosome complex.

To examine whether the predicted SRLase1-EF-Tu·GDP could engage the ribosome, we sought a structure containing ribosome-associated EF-Tu·GDP. Because EF-Tu·GDP normally dissociates rapidly from the ribosome following GTP hydrolysis, such structures are rare. We therefore superimposed the predicted SRLase1-EF-Tu complex onto a cryo-EM structure in which EF-Tu·GDP is retained on the ribosome by the bacterial hibernation factor Balon (PDB code 8RDW) (40). Although *E. coli* does not encode Balon and we do not propose a role for this factor in SRLase1 activity, the structure provides a useful snapshot of a ribosome-associated EF-Tu·GDP state. The EF-Tu molecules aligned closely, positioning the SRLase1 catalytic pocket toward the SRL (Fig. 3H). We obtained a similar result using an independent EF-Tu·GDP-ribosome intermediate captured by time-resolved cryo-EM during translation elongation (PDB code 6WD6) (41), which likewise positioned the catalytic pocket near the SRL despite modest differences in EF-Tu domain orientations (Fig. S13). Together, these observations suggest that the SRLase1-EF-Tu complex can adopt a ribosome-associated configuration that positions the toxin for efficient attack on its target.

These findings identify EF-Tu as an essential host factor for SRLase1-dependent ribosome cleavage. We propose a model in which SRLase1 highjacks EF-Tu and exploits its conformational dynamics and ribosome binding ability to adopt a catalytically active state capable of cleaving the SRL of 23S rRNA (Fig. 4). Consistent with this model, SRLase1 competes with aminoacyl-tRNA for EF-Tu binding and disrupts formation of the EF-Tu·GTP·aa-tRNA ternary complex. This mechanism differs from classical SRL-specific ribotoxins, which directly attack the ribosome, and reveals a host factor-dependent strategy for ribosome targeting by an antibacterial toxin.

**Fig. 4.**
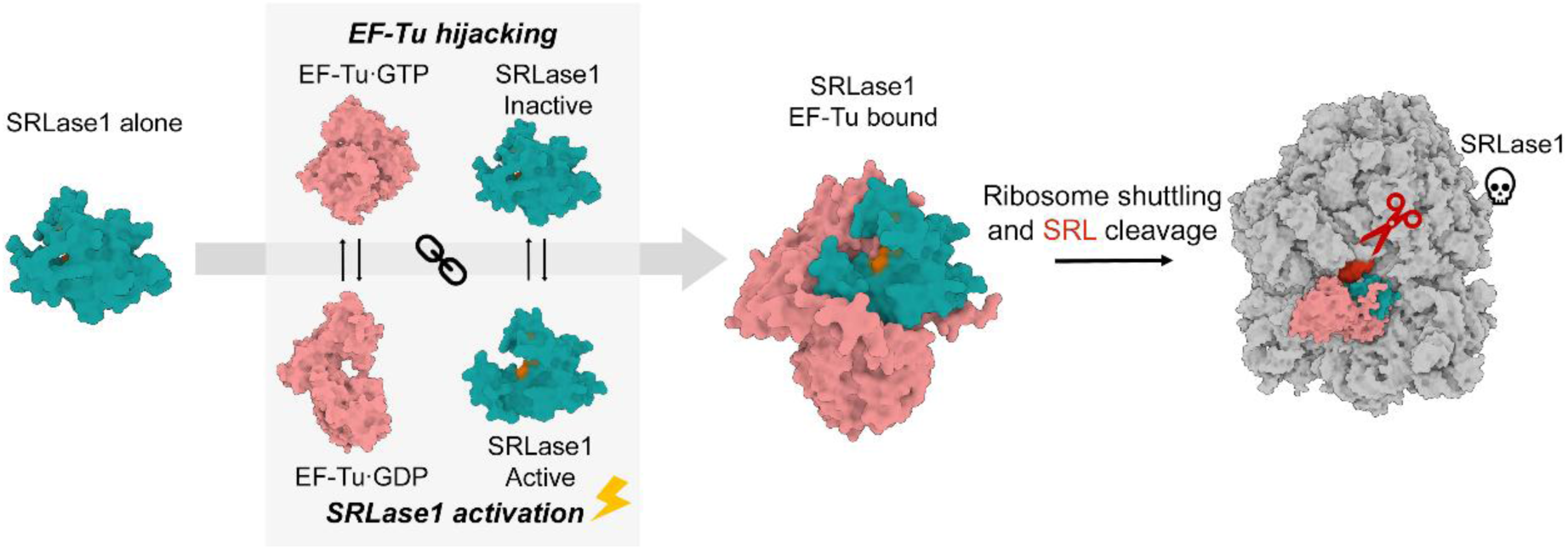
Proposed mechanism of EF-Tu-dependent ribosome cleavage by SRLase1. Unbound and inactive SRLase1 (cyan) interacts with EF-Tu·GDP or EF-Tu·GTP (pink), promoting structural rearrangement of SRLase1 catalytic residues (orange), hijacking the translation factor and cleaving the SRL within ribosomes.

## Discussion

Translation-targeting toxins represent one of the most effective strategies employed during microbial antagonism, yet the molecular mechanisms used to disable protein synthesis continue to expand. Here we identify SRLase1 as the founding member of a family of BECR-like ribotoxins that specifically cleave the SRL of 23S rRNA, a critical component of the GTPase-associated center of the ribosome. Because the SRL coordinates the activity of multiple translational GTPases, cleavage of this single RNA element is expected to disable several steps of protein synthesis, providing an exceptionally efficient route to translational arrest. While numerous antibacterial toxins target translation through cleavage of tRNAs (9–11, 38), mRNAs (6), or modification of translation factors (42) direct attack of the large subunit 23S rRNA is comparatively uncommon. Our findings establish SRLase1 as an antibacterial polymorphic toxin that specifically targets the SRL of 23S rRNA through an EF-Tu-dependent mechanism.

The closest functional parallels to SRLase1 are the classical ribotoxins α-sarcin and restrictocin produced by fungi (43), and the plant ribosome-inactivating protein ricin (34). These toxins also disable translation through attack of the SRL, highlighting the vulnerability of this ribosomal center. The SRL has also emerged as a target in bacterial toxins from TA systems, including VapC20, which cleaves this ribosomal element during stress-associated translational control (13, 14, 44, 45). Thus, SRLase1 joins a small group of toxins that exploit the SRL as a high-value target, but it does so through a distinct molecular strategy. Fungal ribotoxins recognize ribosomes directly without requiring host cofactors (20, 21, 33). In contrast, SRLase1 requires EF-Tu for activity. Moreover, although the SRL sequence is highly conserved across bacterial and eukaryotic ribosomes, SRLase1 did not cleave eukaryotic ribosomes under the conditions tested. These findings indicate that primary sequence alone is insufficient for substrate recognition and suggest that SRLase1 recognizes a specific molecular context created by the bacterial ribosome and its associated translation machinery.

The requirement for EF-Tu distinguishes SRLase1 from previously characterized SRL-targeting toxins and expands the known roles of this translation factor in toxin biology. Several BECR-fold CDI toxins are known to interact with EF-Tu (8, 11, 28, 38). In the best-characterized examples, EF-Tu promotes cleavage of tRNA substrates. For instance, the CDI toxins CdiA-CT^EC869^ and CdiA-CT^NC101^ exploit EF-Tu to recruit tRNAs for cleavage (11, 38), whereas tRNA cleavage by CdiA-CT^Kp342^ is also EF-Tu-dependent, although the underlying mechanism remains unresolved (28). More recently, CdiA-CT^O32:H37^ was proposed to use EF-Tu primarily as an activating or stabilizing factor rather than a substrate-recruitment platform (8). Despite these mechanistic differences, all characterized EF-Tu-associated BECR-fold toxins target tRNAs or other non-ribosomal RNAs. Instead, SRLase1 uses EF-Tu to promote cleavage of the SRL within assembled ribosomes. Because catalytically inactive SRLase1 retained EF-Tu binding but failed to inhibit growth or translation, toxicity is best explained by EF-Tu-dependent SRL cleavage rather than by stoichiometric sequestration of EF-Tu itself.

How EF-Tu promotes SRLase1 activity remains incompletely understood. Several observations suggest that EF-Tu functions as more than a passive binding partner. First, both EF-Tu·GDP and EF-Tu·GTP supported SRLase1-mediated cleavage, arguing against a mechanism restricted to a single nucleotide-bound state. In contrast, kirromycin, which restricts EF-Tu conformational rearrangements, abolished SRL cleavage. Together with the AlphaFold3 prediction that EF-Tu binding remodels the SRLase1 active site, these findings suggest that conformational flexibility is important for formation of a catalytically competent EF-Tu-toxin complex. Structural comparisons further suggest an additional targeting role. Superposition of the predicted SRLase1-EF-Tu complex onto a ribosome-associated EF-Tu·GDP structure positioned the toxin adjacent to the SRL, demonstrating that this ribosome-bound EF-Tu conformation can place SRLase1 within reach of its substrate. We therefore favor a model in which EF-Tu acts both as a ribosome-targeting scaffold and an activating factor, promoting formation of a productive toxin-ribosome complex capable of efficient SRL cleavage (Fig. 4).

The requirement for EF-Tu also raises an evolutionary question: why would a ribosome-targeting toxin evolve dependence on an additional host factor rather than directly recognizing its substrate? One possibility is that SRLase1 evolved from an ancestral EF-Tu-associated toxin that targeted tRNAs, similar to the EF-Tu-dependent CDI toxins discussed above. In this scenario, SRLase1 retained its interaction with EF-Tu while acquiring specificity for the SRL. Rather than becoming dispensable, the pre-existing EF-Tu interaction may have been co-opted as a targeting strategy. Because EF-Tu repeatedly engages ribosomes during translation, retention of this interaction could facilitate delivery of the toxin to its substrate while simultaneously promoting formation of a catalytically competent complex. This interaction may also bias SRLase1 activity toward ribosomes actively involved in translation, where EF-Tu is most frequently recruited. Thus, a host-factor dependency that originally evolved in the context of tRNA cleavage may have been repurposed to enable efficient ribosome targeting. More broadly, this model suggests that toxin evolution can preserve host-factor interactions while redirecting catalytic activity toward new RNA substrates, providing a route for the emergence of novel antibacterial mechanisms.

Collectively, our findings identify SRLase1 as the founding member of a widespread family of SRL-targeting polymorphic toxins and reveal a previously unrecognized mechanism of ribosome inactivation that depends on EF-Tu. Unlike classical ribotoxins, SRLase1 recognizes its substrate only within the context of the translation machinery, demonstrating that host factors can act as critical determinants of toxin specificity. More broadly, these results expand the known strategies used to disable protein synthesis during microbial conflict and suggest that cofactor-assisted ribosome targeting may represent an underappreciated principle among translation-directed toxins.

## Materials and Methods

### Bacterial strains and growth conditions

Bacterial strains used in this study are listed in Table S1. Unless otherwise indicated, bacteria were grown in lysogeny broth (LB; 10 g/L tryptone, 10 g/L NaCl, 5 g/L yeast extract) at 37°C with agitation. When required, media were supplemented with kanamycin at 50 μg/mL, ampicillin at 100 μg/mL, or streptomycin at 50 μg/mL.

### Plasmid construction and mutagenesis

The coding sequence for SRLase1 (FD01848867_02742) was amplified by PCR and cloned into pBRA under control of the arabinose-inducible P_BAD_ promoter (46). SRLi1 (FD01848867_02743) was cloned into pEXT22 under control of the P_TAC_ promoter (47). For complementation assays, *tssL* (FD01848867_02733) and *srli1* (FD01848867_02743) were cloned into pFPV25.1 by replacing the *gfp* mut3 coding sequence with the gene of interest (48). Site-directed mutations were introduced by PCR using primers carrying the desired nucleotide substitutions. *S.* Oslo mutant strains were generated by λ-Red recombineering using the one-step gene inactivation method (49). All plasmids and chromosomal mutations were confirmed by DNA sequencing.

### Interbacterial competition assays

Interbacterial competition assays were performed using *S.* Oslo wild-type or Δ*tssL* strains as attackers and *S.* Oslo Δ*srlase1/srli1*::Km as prey as described previously(50). Overnight cultures were diluted 1:100 in fresh LB and grown to an OD_600nm_ of 0.8 to 1.0. Attacker and prey cultures were adjusted to OD_600nm_ values of 4.0 and 1.0, respectively, and mixed at a 1:1 volume ratio. Five microliters of each mixture were spotted onto 0.22-μm nitrocellulose membranes placed on LB agar plates containing 1.5% agar. After incubation at 37°C for 4 h, membranes were transferred to 1.5-mL tubes containing 1 mL LB and vortexed to resuspend bacteria. Samples were serially diluted and plated on selective media to enumerate surviving prey cells. Prey recovery was calculated as the ratio of output colony-forming units (CFU) to input CFU as described previously (50, 51). For complementation experiments, *S.* Oslo Δ*tssL* attackers carried pFPV25.1-*tssL*, and *S.* Oslo Δ*srlase1/srli1*::Km prey carried pFPV25.1-*srli1*. Non-complemented control strains carried pFPV25.1-*gfp*. Data represents the mean ± SD from six independent experiments. Statistical significance was assessed by one-way analysis of variance followed by Dunnett’s multiple-comparison test, using wild-type competition values normalized to 1.

### *E. coli* toxicity assays

*E. coli* DH5α strains carrying pBRA-SRLase1 variants and, when indicated, pEXT22-SRLi1 were grown overnight in LB containing 0.5% D-glucose as described previously (50). Cultures were adjusted to OD_600nm_ of 1.0, serially diluted fourfold in LB, and spotted onto LB agar containing either 0.5% D-glucose for repression or 0.2% L-arabinose plus 400 μM IPTG for induction of toxin and immunity expression. Plates were supplemented with the appropriate antibiotics and incubated at 37°C. Images were acquired after 20 h.

### Time-lapse microscopy

Time-lapse microscopy was performed using LB agar pads prepared as described previously (52). Briefly, rectangular chambers were cut into double-sided adhesive tape placed on microscopy slides and filled with 1.5% LB agar. *E. coli* DH5α carrying pBRA-SRLase1 was diluted 1:10 in LB containing 0.2% D-glucose and grown to OD_600nm_ 0.4 to 0.6. Cultures were adjusted to OD_600nm_ 1.0 and spotted onto LB agar pads containing either 0.2% D-glucose or 0.2% L-arabinose, with appropriate antibiotics. Images were acquired every 15 min for 16 h using a Leica DMi-8 epifluorescence microscope equipped with a DFC365 FX camera and an HC PL APO 63×/1.4 oil Ph3 objective. Images were analyzed using Fiji (53).

### SOS response assay

The SOS response was monitored using *E. coli* MG1655 carrying the reporter plasmid pSC101-P*_recA_* (54) and pBRA encoding SRLase1_WT_ or SRLase1_H97A_ as described previously (50). Overnight cultures were diluted 1:50 in LB containing 0.5% D-glucose and grown at 37°C to OD_600nm_ 0.4 to 0.6. Cells were harvested and resuspended in AB defined medium containing 0.2% (0.2% (NH4)_2_SO_4_, 0.6% Na_2_HPO_4_, 0.3% KH_2_PO_4_, 0.3% NaCl, 0.1 mM CaCl_2_, 1 mM MgCl_2_, 3 μM FeCl_3_) supplemented with 0.2% sucrose, 0.2% casamino acids, 10 μg/mL thiamine, and 25 μg/mL uracil. Cells were adjusted to OD_600nm_ 1.0 and transferred to black, clear-bottom 96-well plates in a final volume of 200 μL containing either 0.5% D-glucose or 0.2% L-arabinose. GFP fluorescence was monitored for 6 h at 30°C using a SpectraMax Paradigm plate reader.

### RNA extraction and denaturing gel electrophoresis

*E. coli* DH5α strains carrying pBRA-SRLase1 constructs were diluted 1:20 in LB containing 0.5% D-glucose and grown for 1 h at 37°C with agitation at 180 rpm. Cultures were harvested, resuspended in fresh prewarmed LB containing 0.2% L-arabinose, and incubated for an additional 1.5 h to induce toxin expression. Cell pellets corresponding to 3 mL of culture at OD_600nm_ 1.0 were resuspended in 1 mL TRIzol reagent (Invitrogen), and RNA was extracted according to the manufacturer’s instructions. For agarose gel analysis, 5 μg RNA was mixed 1:1 with 2× RNA loading buffer containing 95% formamide, 0.025% SDS, 0.025% bromophenol blue, 0.025% xylene cyanol, and 0.5 mM EDTA. Samples were heated at 70°C for 5 min and resolved on denaturing 1.5% agarose gels containing 0.66 M formaldehyde. Gels were stained in 1× MOPS containing SYBR Gold diluted 1:10,000 and imaged using a ChemiDoc system.

### Primer extension analysis

Primer extension was performed as described previously with modifications (33). Five micrograms of total RNA were used for reverse transcription with the RevertAid Reverse Transcription Kit (Thermo Fisher Scientific). Reactions contained a 5′-FAM-labeled primer complementary to the 3′ region of the 23S rRNA SRL and a nucleotide mixture containing 7 mM ddATP, 10 mM dGTP, 10 mM dTTP, and 10 mM dCTP. Reactions were incubated at 45°C for 1 h. cDNA products were mixed 1:1 with 2× RNA loading buffer, heated at 95°C for 2 min, and resolved on 15% denaturing polyacrylamide gels containing 7 M urea. Electrophoresis was performed in prewarmed 1× TBE buffer at 150 V for 50 min. FAM-labeled oligonucleotides were used as size markers. Gels were imaged using a ChemiDoc system.

### Polysome profiling

*E. coli* BL21(DE3) carrying pBRA encoding FLAG-SRLase1_WT_ or FLAG-SRLase1_H97A_ were diluted 1:50 in 50-mL LB cultures supplemented with streptomycin and 0.5% glucose. Cultures were grown at 37°C with agitation at 180 rpm to OD_600nm_ 0.4 to 0.6, and protein expression was induced with 0.2% arabinose for 1 h at 37°C. After 1 h, cultures were placed on ice and 100 μg/mL chloramphenicol was added and the cultures incubated on ice for 2 min. Cells were then harvested by centrifugation at 6,000 × *g* for 20 min at 4°C. Pellets were stored at −80°C. Cell pellets were thawed on ice and resuspended in 5 mL of Buffer A (10 mM HEPES/KOH pH 7.6, 10 mM MgCl_2_, 100 mM NH_4_Cl, 6 mM β-mercaptoethanol). Cells were lysed at 4°C by sonication at 30% amplitude with 15-s pulses and 45-s pauses for a total sonication time of 3 min. Lysates were clarified by centrifugation at 15,000 × *g* for 15 min at 4°C, and A_260nm_ was measured. Equal A_280nm_ units of each sample were loaded on 10-40% sucrose gradients made with Buffer A. Gradients were spun in a Beckman Coulter Optima XE-90 ultracentrifuge at 31,000 rpm for 3 h at 4°C in an SW 32 Ti swinging bucket rotor. Polysome profiles were analyzed using a BioComp piston gradient fractionator measuring A_260nm_. The A_260nm_ measurements across the gradient were plotted.

### Western blot analysis

All antibodies used are listed in Table S1. For detection of SRLase1 expression in *E. coli*, whole-cell lysates were prepared from cultures grown under repressing or inducing conditions. Samples were mixed with SDS sample buffer, heated at 100°C, and resolved by SDS-PAGE. Proteins were transferred to membranes and probed with antibodies against the epitopes. DnaK was used as a loading control. For *in vitro* translation reactions, translated DHFR-FLAG products were detected by western blot using an anti-FLAG antibody. Anti-EF-Tu was used as a loading control.

### Purification of recombinant SRLase1 proteins for ribosome cleavage assays

*E. coli* BL21(DE3) carrying pRSF-Duet encoding 6×His-SRLase1-Strep and SRLi1 was diluted 1:50 into six 500-mL LB cultures supplemented with kanamycin and 0.5% D-glucose. Cultures were grown at 37°C with agitation at 180 rpm to OD_600nm_ 0.7 to 0.8, and protein expression was induced with 1 mM IPTG for 2 h at 37°C. Cells were harvested by centrifugation at 5,000 × *g* for 20 min at 4°C. Pellets were processed immediately or stored at −80°C. Cell pellets were resuspended in 30 mL buffer A containing 50 mM Tris-HCl, pH 7.5, 350 mM NaCl, and 5% glycerol, supplemented with 5 mg/mL lysozyme. Cells were lysed at 4°C by sonication using 6 to 8 cycles at 70% amplitude with 15-s pulses and 60-s pauses. Lysates were clarified by centrifugation at 40,000 × *g* for 45 min at 4°C, and supernatants were filtered through 0.45-μm low protein-binding PES membranes. Clarified lysate was loaded at 0.5 mL/min onto a 1-mL StrepTrap XT column equilibrated in buffer A. The column was washed with 15 mL buffer A, followed by 10 mL high-salt wash buffer containing 50 mM Tris-HCl, pH 7.5 and 1 M NaCl, and then with an additional 10 mL buffer A. Bound protein was eluted with 6 mL buffer A supplemented with 50 mM biotin.

Biotin-eluted fractions were pooled and subjected to Ni^2+^-affinity chromatography using a 1-mL HisTrap HP column. The column was equilibrated in binding buffer containing 50 mM Tris-HCl, pH 7.5, 350 mM NaCl, 5% glycerol, and 5 mM β-mercaptoethanol. Samples were loaded at 0.5 mL/min, and the column was washed with 10 CV of binding buffer. To remove co-purifying proteins and refold SRLase1 on-column, bound protein was denatured using a linear 0 to 100% gradient of denaturing buffer, consisting of binding buffer supplemented with 6 M guanidine-HCl, over 20 CV at 0.8 mL/min. The column was then washed with 8 CV of 100% denaturing buffer. Refolding was performed by applying a reverse 100 to 0% denaturing buffer gradient over 20 CV at 0.1 mL/min, followed by 8 CV of binding buffer. Refolded protein was eluted using a linear 0 to 100% gradient of elution buffer, consisting of binding buffer supplemented with 500 mM imidazole, over 20 CV, followed by 8 CV of 100% elution buffer. Protein-containing fractions were pooled and buffer exchanged into storage buffer containing 20 mM Tris-HCl, pH 7.5, 100 mM NaCl, and 5% glycerol using a 10-kDa molecular weight cutoff Amicon centrifugal filter.

### Purification of SRLase1 proteins for microscale thermophoresis

SRLase1 proteins used for MST were purified from *E. coli* BL21(DE3) carrying pRSF-Duet-SRLase1-Strep:SRLi1 constructs using the StrepTrap XT purification procedure described above, with the following modifications. For SRLase1_H97A_-Strep, the column was washed with 5 CV of buffer A supplemented with 3 M urea to remove co-purifying EF-Tu, followed by 15 CV of buffer A. Protein was eluted with 3 CV of buffer A supplemented with 50 mM biotin, buffer exchanged using a 10-kDa Amicon, and stored in storage buffer. For SRLase1_WT_, purification under native conditions yielded a stronger SRLase1–SRLi1 complex. Therefore, the complex was denatured overnight in buffer A containing 6 M urea, reapplied to the StrepTrap XT column, and eluted under denaturing conditions to separate SRLase1 from SRLi1. Purified SRLase1 was refolded using a stepwise dilution procedure as done previously with adaptations (55). Briefly, eluted protein was diluted twofold into ice-cold storage buffer and incubated on ice for 10 min. This dilution step was repeated, and the resulting 4-mL suspension was concentrated to 500 μL using a 10-kDa Amicon. The concentrated protein was diluted into 5 mL ice-cold buffer A. Concentration and dilution steps were repeated twice, followed by final buffer exchange into 30 mL storage buffer.

### Purification and nucleotide exchange of EF-Tu

6×His-tagged EF-Tu was purified as described previously with modifications (56). *E. coli* BL21(DE3) carrying pCA24N-*tufA* was grown in 1 L terrific broth (24 g/L yeast extract, 20 g/L tryptone, 4 mL/L glycerol, 0.017 M KH_2_PO_4_, 0.072 M K_2_HPO_4_) at 37°C to OD_600nm_ 0.6. EF-Tu expression was induced with 1 mM IPTG for 3 h at 37°C. Cells were harvested by centrifugation at 10,000 × *g* for 10 min and stored at −80°C. Cell pellets were resuspended in lysis buffer containing 50 mM Tris-HCl pH 7.6, 60 mM NH4CI, 7 mM MgCl2, 7 mM β-Me, 15% glycerol, 10 µM GDP, 10 mM imidazole. Cells were lysed by sonication, and insoluble material was removed by centrifugation at 28,000 × *g* for 15 min. The clarified lysate was applied to a 1-mL HisTrap HP column equilibrated in lysis buffer. The column was washed with 10 CV of lysis buffer, followed by 10 CV of wash buffer 1 containing 50 mM Tris-HCl pH 8.0, 60 mM NH_4_Cl, 7 mM MgCl_2_, 7 mM β-Me, 15% glycerol, 10 µM GDP, 10 mM imidazole, 300 mM KCl. The column was then washed with 10 CV of wash buffer 2 containing 50 mM Tris-HCl pH 7.0, 60 mM NH_4_Cl, 7 mM MgCl_2_, 7 mM β-Me, 15% glycerol, 10 µM GDP, 10 mM imidazole, 300 mM KCl. EF-Tu was eluted stepwise with elution buffer containing 50 mM Tris-HCl pH 7.6, 60 mM NH_4_Cl, 7 mM MgCl_2_, 7 mM β-Me, 15% glycerol, 10 µM GDP, and increasing concentrations of imidazole at 25, 50, 75, 100, 125, 250, and 500 mM. For nucleotide exchange, eluted EF-Tu was buffer exchanged into 25 mM Tris-HCl pH 7.5, 50 mM NH_4_Cl, 10 mM EDTA using a 10-kDa Amicon filter and incubated at 37°C for 10 min. Apo EF-Tu and released GDP were separated by size-exclusion chromatography on a HiLoad 16/600 Superdex 200 pg column equilibrated in 25 mM Tris-HCl, pH 7.5, and 50 mM NH_4_Cl. GTP or GDP was added immediately after purification to a final concentration of 20 μM. EF-Tu was stored in 50% glycerol at −20°C.

### Microscale thermophoresis binding assays

MST was used to measure SRLase1 binding to EF-Tu. 6×His-tagged EF-Tu·GTP at 150 nM was labeled with 50 nM RED-tris-NTA dye, second generation (NanoTemper Technologies), in binding buffer containing 1× PBS, 50 mM NaCl, 30 mM KCl, and 0.01% Tween 20. Labeling reactions were incubated at room temperature (RT) for 30 min and centrifuged at 15,000 × *g* for 10 min at 4°C as described previously (56). Labeled EF-Tu was mixed 1:1 with serial dilutions of SRLase1-Strep and incubated at RT for 2 h. Measurements were performed using a Monolith NT.115 instrument. Changes in fluorescence were plotted as a function of SRLase1 concentration and fitted to the hyperbolic function to estimate apparent binding constants:

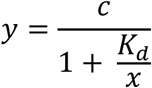

where *y* is the measured fluorescent signal, *c* is the difference between free target and bound target-ligand complex, K_d_ is the dissociation constant (ligand concentration at which half of all target molecules are bound), and x is the concentration of free ligand in the assay.

### Ribosome cleavage assays

Purified *E. coli* 70S ribosomes (New England Biolabs, #P0763S) were incubated at a final concentration of 200 nM with 20 nM SRLase1_WT_ or SRLase1_H97A_, unless otherwise indicated, in the presence or absence of 200 nM EF-Tu. Reactions were performed in cleavage buffer containing 25 mM Tris-HCl, pH 7.5, 50 mM NaCl, 30 mM KCl, 8 mM MgCl_2_, and 5 mM β-mercaptoethanol. Reactions were incubated at 37°C for 15 min. Where indicated, EF-Tu was preincubated with 1 mM kirromycin (BioViotica, BVT-0157-M001) for 5 min at 37°C before addition of SRLase1 and ribosomes. RNA was extracted with TRIzol and resolved on 5% denaturing PAGE containing 7 M urea in 0.5× TBE. Gels were stained for 30 min with SYBR Gold diluted 1:10,000 in TBE and imaged using a ChemiDoc system.

### *In vitro* translation assays

*In vitro* translation reactions were performed using the PURExpress kit (New England Biolabs) according to the manufacturer’s instructions. Reactions were assembled with the pM2-DHFR-FLAG reporter plasmid and 900 nM SRLase1_WT_ or 900 nM SRLase1_H97A_. Reactions were incubated at 37°C for 2 h. Half of each reaction was mixed with SDS sample buffer, heated at 100°C for 5 min, and analyzed by SDS-PAGE followed by western blot. The remaining reaction volume was mixed with TRIzol for RNA extraction. *In vitro* translation in eukaryotic system was performed using the 1-Step Human Coupled IVT Kit (ThermoFisher, #88881) according to manufactures’ instructions. Reactions were assembled containing 400 nM SRLase1_WT_ or 400 nM SRLase1_H97A_, with and without 400 nM of EF-Tu·GDP. Reactions were incubated at 37°C for 2.5 h. Half of each reaction was mixed with SDS sample buffer and analyzed by western blot. The remaining reaction volume was mixed with TRIzol for RNA analysis.

### Acidic extraction of total tRNA

Total tRNA was extracted from *E. coli* BW25113 Δ*rnr* cells under acidic conditions as described previously with modifications (57). Overnight cultures were diluted 1:25 in LB and grown for 3 h. Cells were rapidly chilled on ice, harvested by centrifugation, and resuspended in 300 μL extraction buffer containing 0.3 M sodium acetate, pH 4.5, and 10 mM EDTA. Samples were extracted twice with 300 μL phenol, and RNA was precipitated with 750 μL ice-cold 100% ethanol for 1.5 h on ice. RNA was collected by centrifugation at 12,000 × *g* for 15 min and washed twice with 150 μL 1 M NaCl, pH 4.5, to remove larger ribosomal RNA species and proteins. After each wash, samples were centrifuged at 12,000 × *g* for 30 min, and supernatants were collected. Combined supernatants were ethanol precipitated, centrifuged at 12,000 × *g* for 15 min, and resuspended in 10 mM potassium acetate to a final volume of 20 μL. Samples were stored at −80°C or used immediately.

### Native gel-shift assays

EF-Tu·GTP·aa-tRNA ternary complexes were assembled as described previously with modifications (58, 59). EF-Tu at 5 μM was incubated with 10 mM GTP in buffer 1 containing 70 mM HEPES-KOH, pH 7.6, 52 mM NH_4_OAc, 8 mM Mg(OAc)_2_, 230 mM KCl, and 0.8 mM DTT. Reactions were performed in a volume of 2.5 μL and incubated at 37°C for 5 min. Aminoacyl-tRNA was then added at 1.3 μg in 1.5 μL of 10 mM potassium acetate, together with 1 μL buffer 2 containing 150 mM HEPES-KOH, pH 7.6, 195 mM NH_4_OAc, and 30 mM Mg(OAc)_2._ Reactions were incubated at 37°C for 10 min. To test whether SRLase1 disrupts preassembled ternary complexes, SRLase1 was added after ternary complex formation at the indicated volumes of 2.5, 5, or 10 μL from a 7.7 μM stock prepared in 20 mM Tris-HCl, pH 7.5, 100 mM NaCl, and 5% glycerol. Reactions were incubated for 5 min at 37°C. To test whether SRLase1 prevents ternary complex formation, EF-Tu and SRLase1 were preincubated for 5 min at 37°C before addition of aminoacyl-tRNA. Samples were mixed with 1.5 μL of 50% glycerol containing bromophenol blue. Reactions containing GTP were supplemented to a final concentration of 1 mM GTP immediately before electrophoresis. Complexes were resolved on 5% native PAGE prepared with a 19:1 acrylamide:bis-acrylamide ratio in 50 mM Tris-HCl, pH 6.8, 65 mM NH_4_OAc, and 10 mM Mg(OAc)_2_. Electrophoresis was performed at 4°C for 1.5 h at 110 V. Gels were stained by silver staining.

### Bioinformatic analysis

Sequences from the SRLase1 group identified by Nicastro et al. (2026) were used as queries for PSI-BLAST searches against the UniRef50 database. Searches were performed for three iterations. Resulting alignments were manually curated to retain homologs preserving the predicted secondary-structure elements and conserved residues characteristic of the BECR-like fold, including an N-terminal α-helix and at least three β-strands. Sequence logos were generated using WebLogo (60). Final alignment models were used for searches against the NCBI nonredundant protein database to identify homologs for taxonomic distribution, with 100% redundant sequences removed. Genomic neighborhoods were collected for homologs using ROTIFER (https://github.com/leepusp/rotifer). Domain identification and annotation were performed using HMMER tools (61–63), together with Pfam models (64). Taxonomic distribution was visualized using AnnoTree (65).

### Statistical analysis

All statistical analyses were performed in GraphPad Prism 5. Statistical analyses were performed as indicated in the figure legends. For interbacterial competition assays, data represents the mean ± SD from six independent experiments. Comparisons were performed by one-way analysis of variance followed by Dunnett’s multiple-comparison test.

## Acknowledgements

This research was supported by startup funds from UT Austin College of Natural Sciences to E.B-S, Sao Paulo Research Foundation grant # 2021/10577-0 to R.F.S. This work was supported by the National Institutes of Health (NIH) R35 GM127127 to A.J., R01 GM121650 to K.K., R35 GM156629 to C.M.D. and NIH T32 AI106699 to T.M.B. C.M.D. is a Burroughs Wellcome Fund Investigator in the Pathogenesis of Infectious Diseases. The authors acknowledge the Texas Advanced Computing Center (TACC) at The University of Texas at Austin for providing computational resources that have contributed to the research results reported within this paper (http://www.tacc.utexas.edu). Additional support was provided in part by an appointment to the National Library of Medicine Research Participation Program administered by the Oak Ridge Institute for Science and Education (ORISE) though an interagency agreement between the U.S. Department of Energy (DOE) and the National Library of Medicine. ORISE is managed by ORAU under DOE contract number DE-SC0014664. All opinions expressed in this paper are the author’s and do not necessarily reflect the policies and views of NIH, NLM, DOE, or ORAU/ORISE. This work is supported by the funds of the Intramural Research Program of National Library of Medicine at the National Institutes of Health, USA (G.G.N).

## Author Contributions

J.T.H. and E.B.S designed the study. J.T.H., T.M.B. and A.V. conducted experiments. K.P., G.G.N. and R.F.S. performed bioinformatic analysis. L.A., A.J. K.K. and C.M.D. contributed with scientific insights and discussions. J.T.H. and E.B.S. wrote the manuscript with input from all authors.

