## Supplementary material for "An RNase Toxin Hijacks Elongation Factor-Tu to Cleave the Ribosomal Sarcin-Ricin Loop": SI Appendix

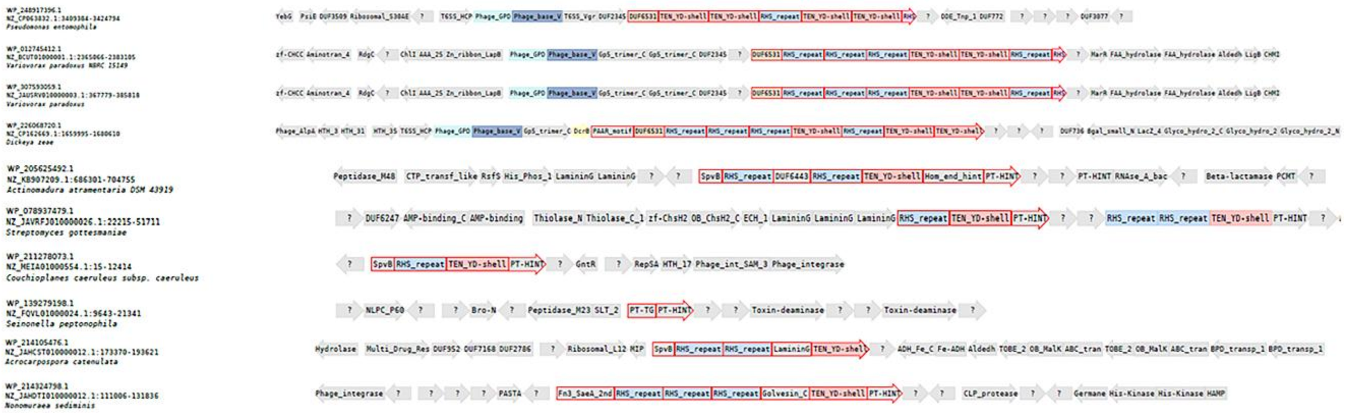


**Fig. S1. Genomic context of SRLase1 homologs**. SRLase1 homologs (red border) and neighbors with identified domains annotated.

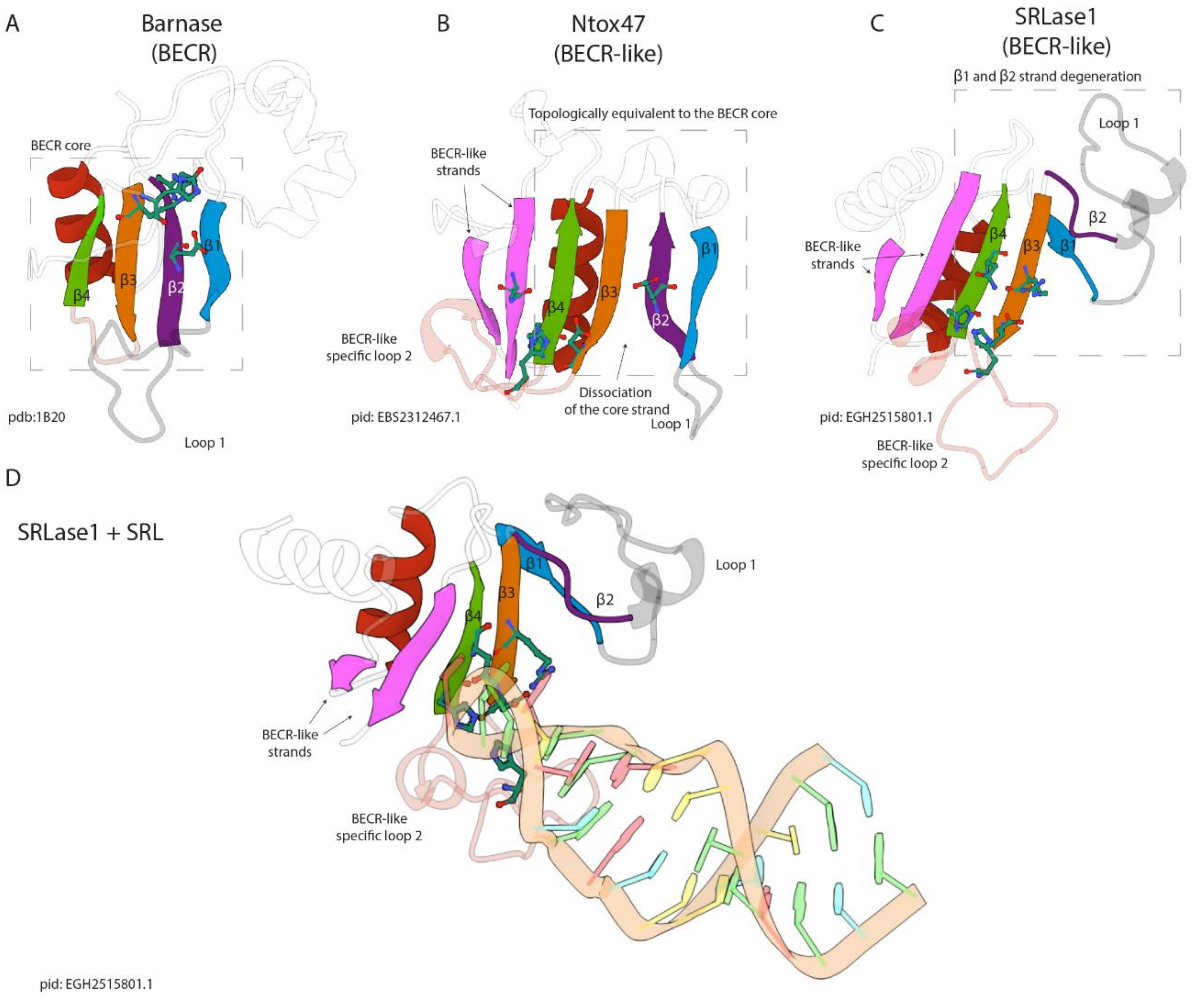


**Fig. S2. Comparison of strand organization in classical BECR and BECR-like RNases.** (a) Structure of the canonical BECR-fold Barnase, illustrating the conserved architecture consisting of an N-terminal α-helix followed by a four-stranded β-meander (β1-β4) that presents the catalytic surface. (b-c) AlphaFold3-predicted structures of representative BECR-like proteins Ntox47 and SRLase1. The conserved β-sheet core of the BECR-like superfamily is retained but differs from canonical BECR proteins by the addition of a C-terminal β-strand (β5). In some members, an additional N-terminal β-strand packs parallel to β5, further extending the sheet. Strands topologically equivalent to β1 and β2 of classical BECR proteins display varying degrees of degeneration and dissociation from the core sheet, whereas the remaining sheet architecture is strongly conserved across the superfamily. (d) AlphaFold3 model of SRLase1 bound to the sarcin-ricin loop (SRL). Variable regions corresponding to Loop1 and Loop2 are highlighted.

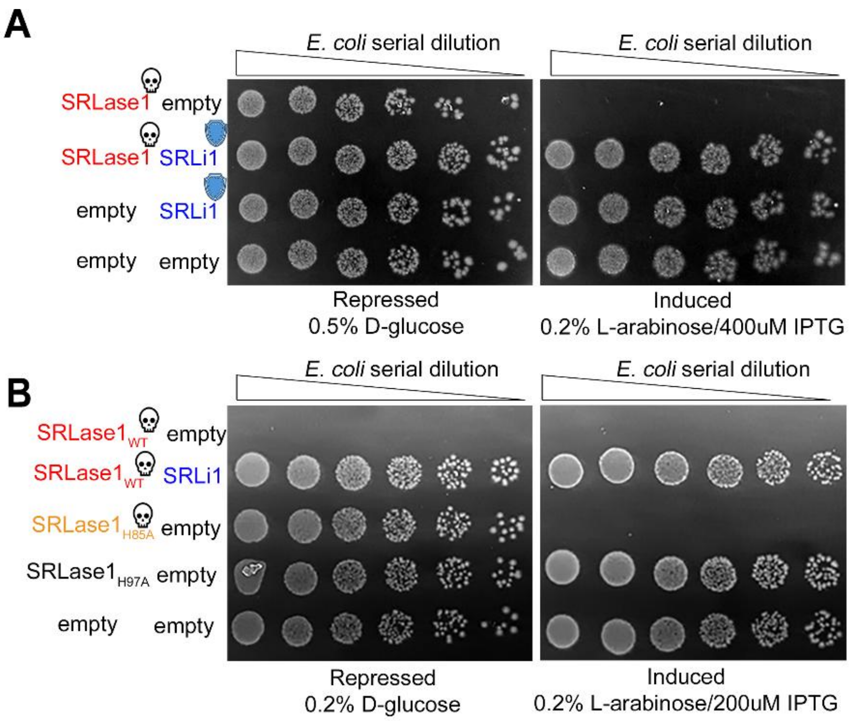


**Fig. S3. SRLase1 toxicity under reduced repression.** *E. coli* serial dilution assay of cells expressing pBRA-SRLase1 variants and pEXT22-SRLi1. **(a)** Under standard repressing conditions (0.5% glucose), SRLase1$_{H85A}$ displayed toxicity comparable to wild-type SRLase1. However, under reduced repressing conditions (0.2% glucose) **(b)**, the H85A mutant showed a reproducible decrease in toxicity. Cells were grown under repressing (0.2% glucose) or inducing conditions (0.2% L-arabinose).

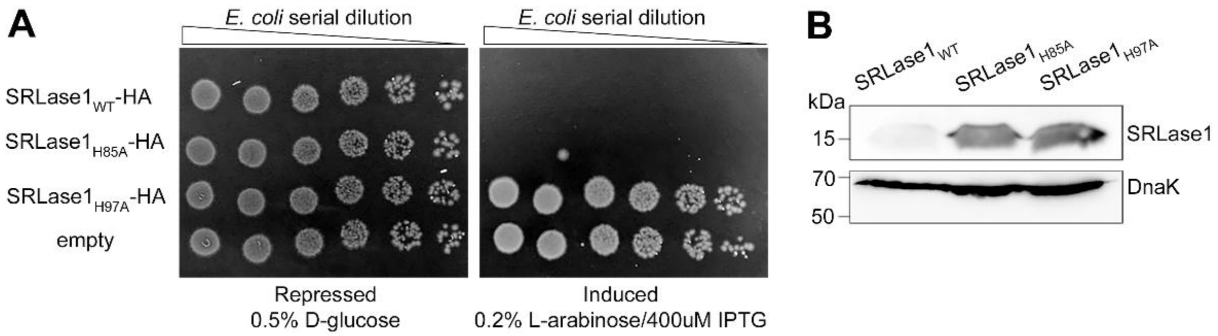


**Fig S4. SRLase1 mutants are expressed at comparable levels. (a)** *E. coli* serial dilution toxicity assay of cells expressing HA-tagged wild-type or mutant SRLase1 proteins, as indicated. **(b)** Western blot of whole-cell lysates following 2 h induction with 1% L-arabinose. Comparable accumulation of wild-type and mutants was detected using the C-terminal HA tag. Anti-DNAk was used as loading control.

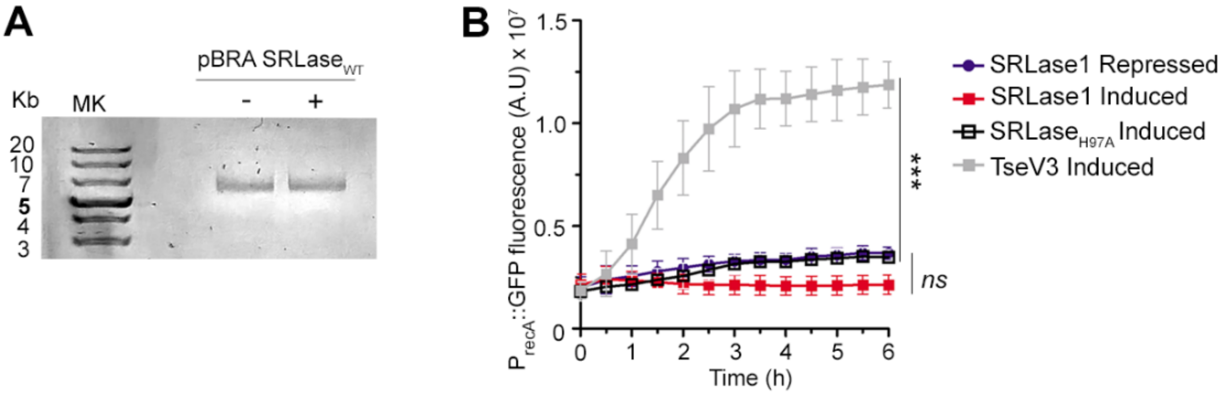


**Fig S5. SRLase1 does not induce DNA damage. (a)** Plasmid DNA integrity following SRLase1 expression. Plasmids were isolated from *E. coli* grown for 1 h under repressing (0.5% D-glucose) or inducing (0.2% L-arabinose) conditions and analyzed by agarose gel electrophoresis. **(b)** SOS response assay using *E. coli* carrying the reporter plasmid pSC101-$P_{recA}$::GFP (Ronen et al., 2002). Cells expressing SRLase1 or the DNase toxin TseV3 (positive control; Hespanhol et al., 2022) were grown in AB medium under repressing or inducing conditions. Fluorescence values are shown as mean ± SD from three independent experiments. Statistical significance was assessed by one-way ANOVA followed by Dunnett's multiple-comparison test. ns, not significant; ***P* < 0.001.

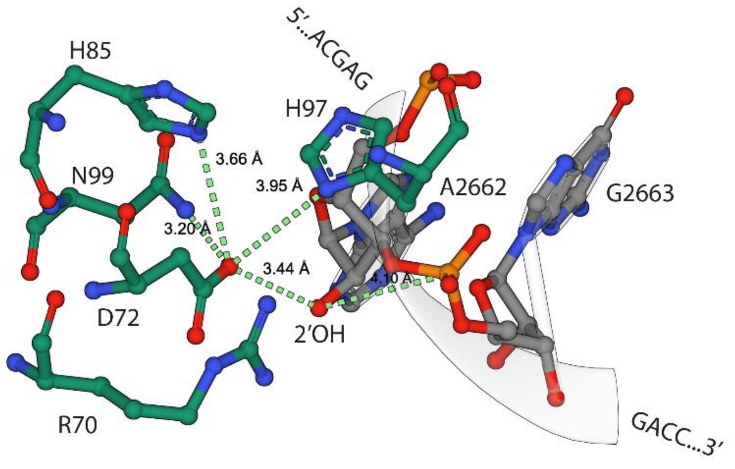


**Fig. S6. Predicted conserved residues involved in SRL cleavage.** AlphaFold3 model of SRLase1 bound to the sarcin–ricin loop (SRL), showing positioning of the substrate within the catalytic pocket. Conserved residues are arranged in a manner consistent with the chemistry of metal-independent RNA cleavage.

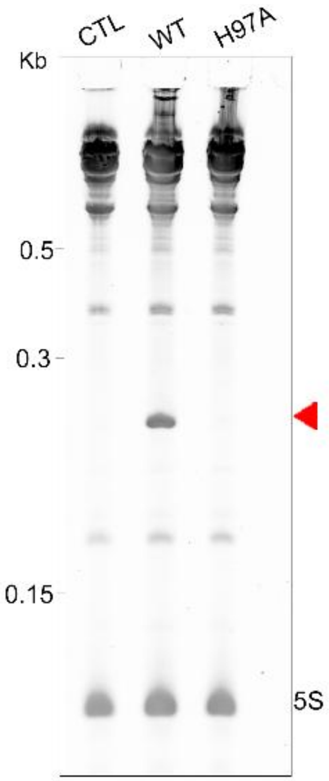


**Fig. S7. SRLse1 cleaves the SRL during *in vitro* translation.** RNA extracted from *in vitro* translation reactions (Fig. 2G) and analyzed via denaturing PAGE. The SRL cleavage fragment generated by SRLase1 activity is indicated (red triangle).

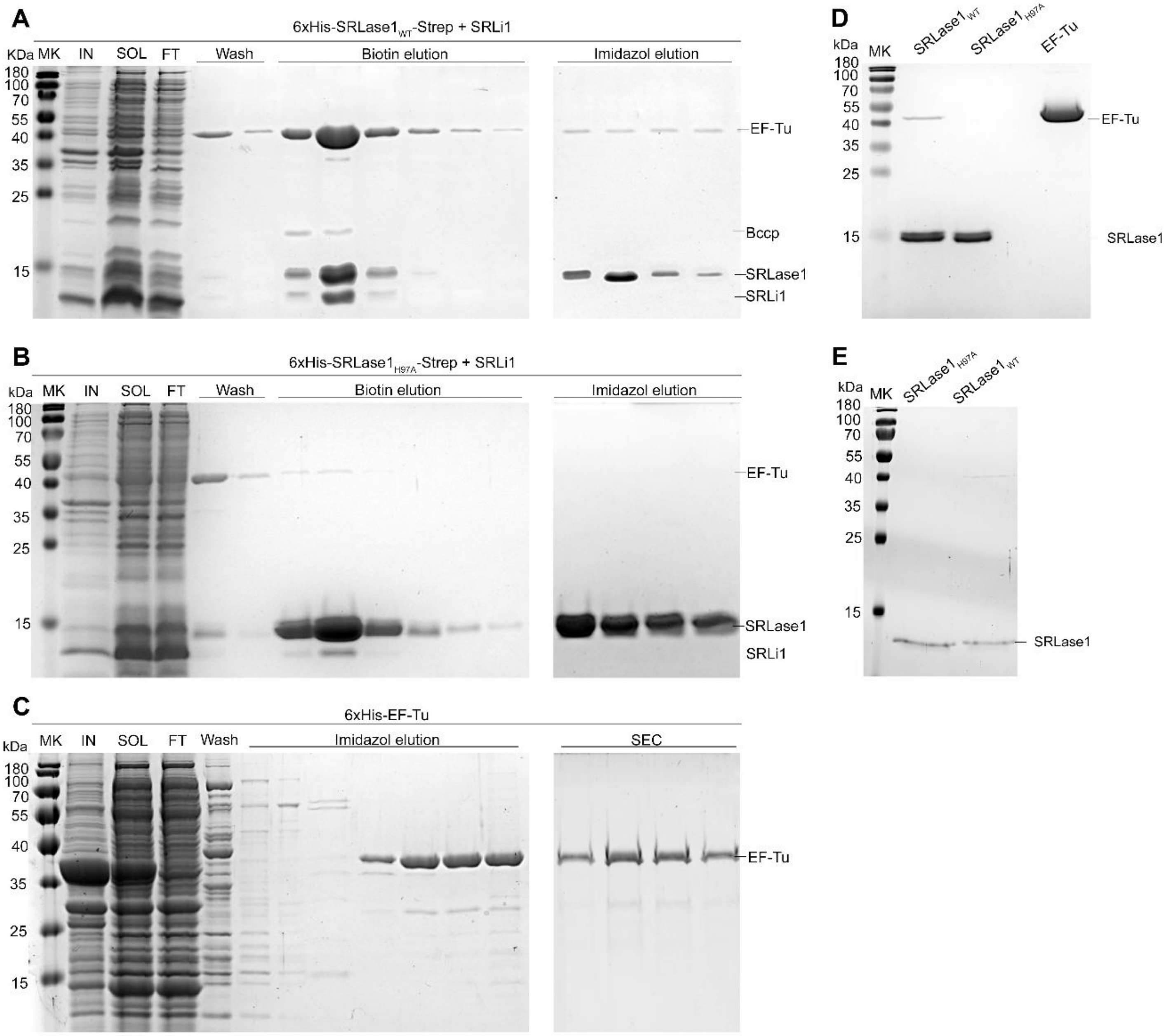


**Fig S8. Purification of recombinant SRLase1 and EF-Tu proteins. (a-b)** SDS–PAGE of recombinant **(a)** SRLase1$_{WT}$ or **(b)** SRLase$_{H97A}$ during the two-step purification procedure used for ribosome cleavage assays. Shown are samples from the first Strep affinity purification, including the insoluble fraction (**I**), soluble fraction (SOL), flowthrough (FT), wash (W), and biotin-eluted fractions. The biotin-eluted material was subjected to HisTrap HP chromatography under denaturing/refolding conditions, followed by imidazole elution. **(c)** Affinity purification and size exclusion chromatography (SEC) of recombinant EF-Tu. **(d)** Purified and refolded His-SRLase1$_{WT}$-Strep and His-SRLase$_{H97A}$-Strep, and EF-Tu proteins used in ribosome cleavage assays. (**e**) Purified and refolded proteins SRLase1$_{WT}$-Strep and SRLase$_{H97A}$-Strep for MST binding assays.

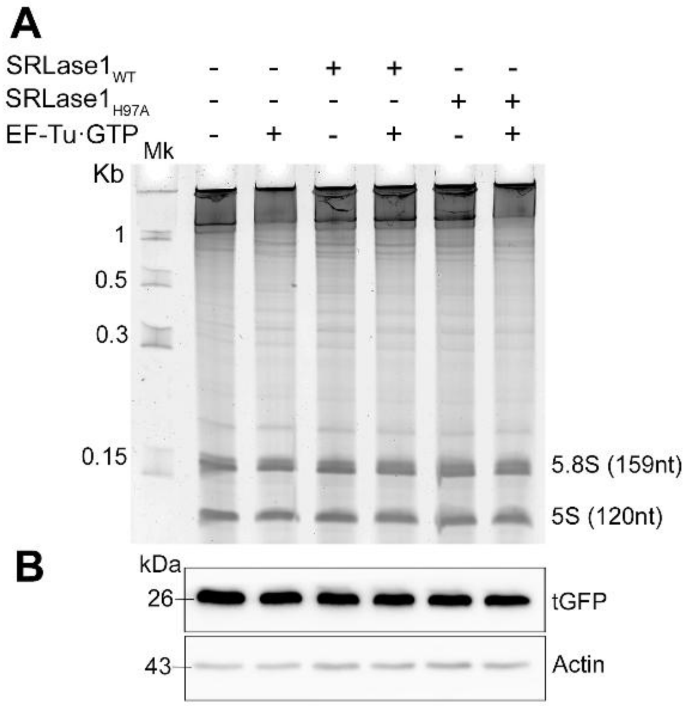


**Fig. S9. SRLase1 does not target eukaryotic ribosomes *in vitro*.** *In vitro* translation assay performed in the of recombinant SRLase1$_{WT}$, or SRLase1$_{H97A}$, with and without EF-Tu·GTP. **(a)** Denaturing PAGE of the RNA extracted from the reaction. **(b)** Translation output of a GFP reporter monitored by western blot. Anti-actin was used as loading control.

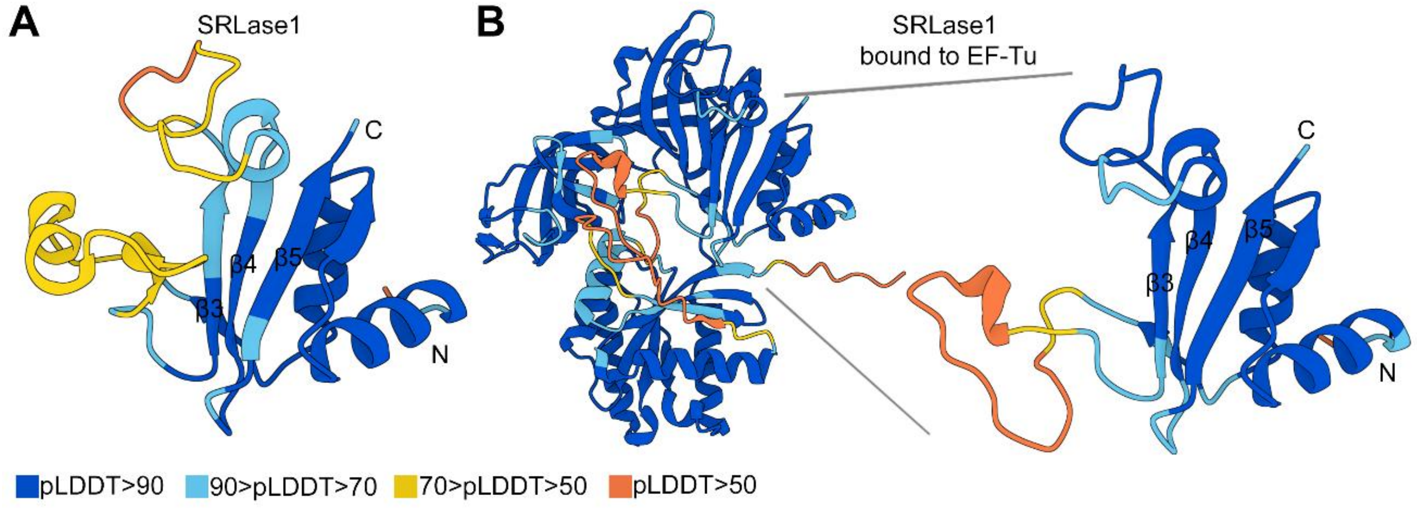


**Fig S10. AlphaFold3 confidence scores for the SRLse1-EF-Tu complex. (a)** AlphaFold3 prediction of SRLase1 alone. **(b)** AlphaFold3 prediction of the SRLase1-EF-Tu complex. Structures are colored according to pLDDT, a per-residue confidence score ranging from 0 to 100, with higher values indicating increased prediction confidence. EF-Tu UniProtKB accession is P0CE47.1.

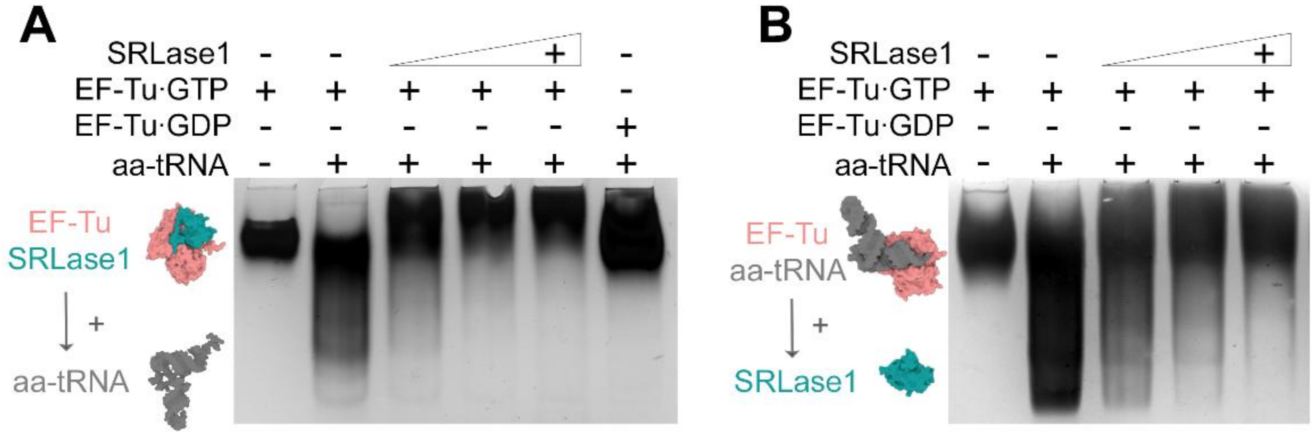


**Fig. S11. SRLase1 disrupts EF-Tu·GTP·aa-tRNA ternary complex formation.** Native gel-shift assays showing the effect of SRLase1 on EF-Tu·GTP·aa-tRNA ternary complexes. Increasing amounts of SRLase1 were added as indicated. **(a)** EF-Tu and SRLase1 were preincubated before addition of aa-tRNA. **(b)** The EF-Tu·GTP·aa-tRNA ternary complex was assembled prior to addition of SRLase1.

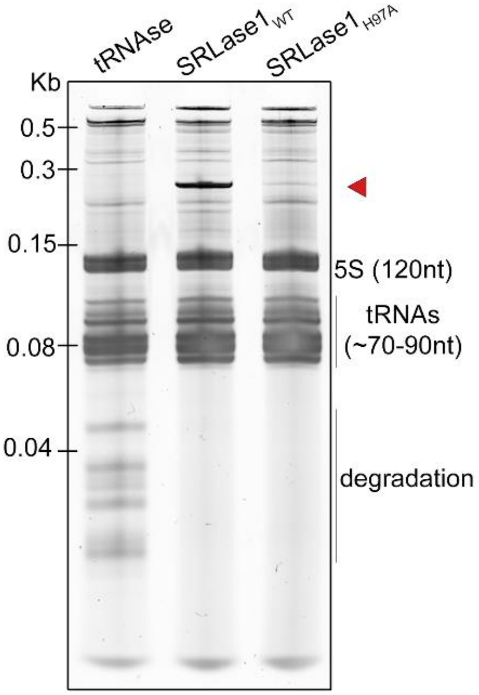


**Fig. S12. SRLase1 does not produce detectable tRNA cleavage products.** Analysis of total RNA extracted from *E. coli* following expression of the indicated SRLase1 variants. A characterized tRNase toxin was included as a positive control to illustrate the expected RNA cleavage pattern.

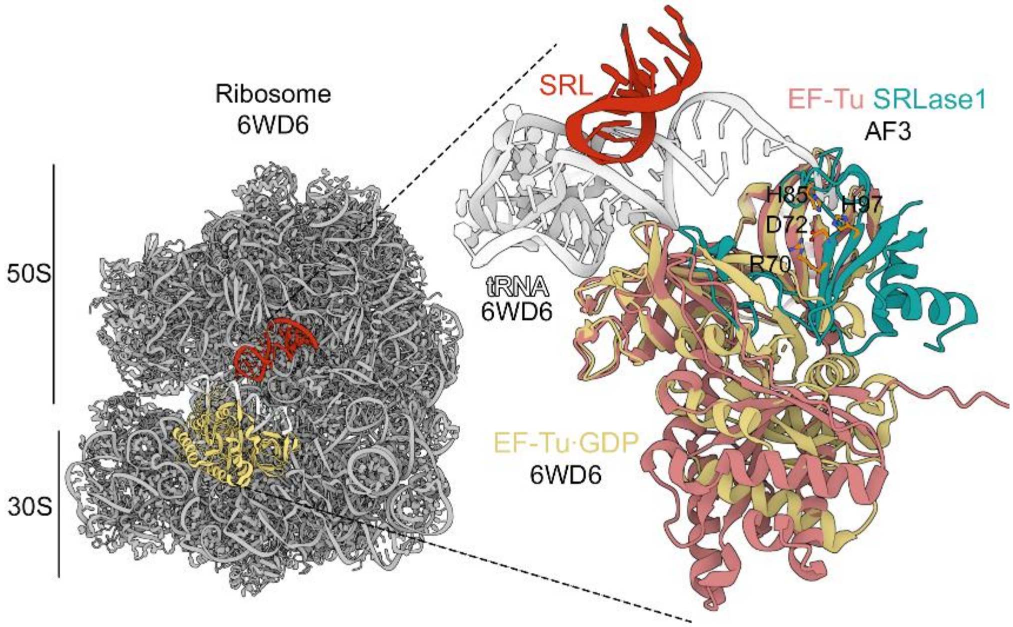


**Fig. S13. Comparison of the EF-Tu–SRLase1 model with ribosome-bound EF-Tu·GDP.** Structural overlay of the AlphaFold3 (AF3) model of the EF-Tu–SRLase1 complex with EF-Tu–GDP from the ribosome-bound structure (PDB 6WD6).

### Tables

**Table S1.** Table S1. List of strains, plasmids, antibodies and primers used in this study.

| Strains | Description | Source |
|---|---|---|
| *Salmonella* enterica Oslo | *S.* Oslo (01848867) | [1] |
| *Escherichia coli* DH5α | Cloning and toxicity assays | Lab stock |
| *Escherichia coli* BL21(DE3) | Recombinant protein expression | Lab stock |
| *Salmonella* enterica Oslo Δ*tssL* | *S.* Olso Δ*tssL* | This study |
| *Salmonella* enterica Oslo Δ*srlase1/srli1* | *S.* Oslo Δ*srlase1/srli1*:: KmR | This study |
| **Plasmids** | **Description** | **Reference** |
| pBRA | Derivative of pBAD24, $P_{BAD}$ promoter, mob, pBBR1 ori, $Sp^R$ | [2] |
| pBRA SRLase1$_{WT}$ | pBRA expressing SRLase1$_{WT}$ (FD01848867_02742) to *E. coli* toxicity assay | This study |
| pBRA SRLase$_{H85A}$ | pBRA expressing SRLase1$_{H85A}$ (FD01848867_02742) to *E. coli* toxicity assay | This study |
| pBRA SRLase$_{H97A}$ | pBRA expressing SRLase1$_{H97A}$ (FD01848867_02742) to *E. coli* toxicity assay | This study |
| pBRA SRLase1$_{WT}$-HA | pBRA expressing SRLase1$_{WT}$ (FD01848867_02742) to *E. coli* toxicity assay | This study |
| pBRA SRLase$_{H85A}$-HA | pBRA expressing SRLase1$_{H85A}$ (FD01848867_02742) to *E. coli* toxicity assay | This study |
| pBRA SRLase$_{H97A}$-HA | pBRA expressing SRLase1$_{H97A}$ (FD01848867_02742) to *E. coli* toxicity assay | This study |
| pBRA FLAG-SRLase1$_{WT}$ | pBRA expressing SRLase1$_{WT}$ (FD01848867_02742) for polysome | This study |
| pBRA FLAG-SRLase$_{H97A}$ | pBRA expressing SRLase1$_{H97A}$ (FD01848867_02742) for polysome | This study |
| pEXT22 | $P_{TAC}$ promoter, R100 ori, $Km^R$ | [3] |
| pEXT22 SRLi1 | pEXT22 expressing SRLi1 (FD01848867_02743) for *E. coli* toxicity assay | This study |
| pFPV25.1 | PrpsM promoter, GFP mut3, pBR322 ori, AmpR | [4] |
| pFPV25.1 TssL | pFPV25.1 expressing TssL for bacterial competition | This study |
| pFPV25.1 SRLi1 | pFPV25.1 expressing SRLi1 for bacterial competition | This study |
| pKD4 | Gene deletion by allelic exchange/PCR template (KmR) | [5] |
| pKD46 | Gene deletion by allelic exchange/Lambda Red genes | [5] |
| pCP20 | Gene deletion by allelic exchange/FRT Recombinase | [5] |
| pSC101-$P_{recA}$::GFP | $P_{recA}$::GFP, pSC101 ori, $Km^R$ | [6] |
| pRSF-Duet | PT7, RSF1030 ori, lacI, KmR | Novagen cat #71341-3 |
| pRSF-Duet SRLase1$_{WT}$+SRLi1 | pRSF::6xHis-SRLase1$_{WT}$-Strep::TsiV3 | This study |
| pRSF-Duet SRLase1$_{H97A}$+SRLi1 | pRSF::6xHis-SRLase1$_{H97A}$-Strep::TsiV3 | This study |
| pCA24N His-*tufA* | $P_{T5}$ promoter IPTG-inducible expression of 6xHis-EF-Tu; pMB1 ori, $Cm^R$ | [7] |
| pM2-DHFR | Expresses DHFR-FLAG, T7 promoter; AmpR | [7] |
| **Antibodies** | **Description** | **Source** |
| Strep-Tactin HRP conjugate | Western blot | Fisher Scientific Cat# NC9523094 |
| Mouse anti-EF-Tu | Western blot | Abcam Cat#ab90813 |
| Rabbit anti-HA | Western blot | Sigma Cat#H6908 |
| Rabbit anti-FLAG | Western blot | Sigma Cat#F2555 |
| **Primers** | **Sequence** | **Purpose** |
| EBS 537-F NcoI | AATTCCATGGGTGCTGGATGTAATGCCCAAC | Clone SRLase1WT (FD01848867_02742) in pBRA |
| EBS 538-R SalI | TATAGTCGACCTAAGGAACGATAATTATCCCAG | Clone SRLase1WT (FD01848867_02742) in pBRA |
| EBS 539-F BamHI | AATAGGATCCAGGAGGAATTCACCATGAGAAATACAATAGATAATAGG | Clone SRLi1 (FD01848867_02743) in pEXT22 |
| EBS 540-R HindIII | ATATAAGCTTTTATAATTTATCCATCCACGA | Clone SRLi1 (FD01848867_02743) in pEXT22 |
| EBS 621-F | CATTATCTAATGTTGTT**GCG**AATTCTAAAAATC | Point mutation H85A in SRLase1 |
| EBS 622-R | GATTTTTAGAATT**CGC**AACAACATTAGATAATG | Point mutation H85A in SRLase1 |
| EBS 623-F | GATAATAATCCG**GCG**TATAATATTATCTTTGG | Point mutation H97A in SRLase1 |
| EBS 624-R | CCAAAGATAATATTATA**CGC**CGGATTATTATC | Point mutation H97A in SRLase1 |
| EBS 629-F XbaI | AAATCTAGAAAGAAGGAGATATACATATGAGAAATACAATAGATAATAGG | Clone SRLi1 (FD01848867_02743) in pFPV25.1 |
| EBS 630-R HindIII | TTTTAAGCTTTTATAATTTATCCATCCACGA | Clone SRLi1 (FD01848867_02743) in pFPV25.1 |
| EBS 729-F XbaI | AAATCTAGATTTAAGAAGGAGATATACATATGACAGACAGTACCCTGAC | Clone *tssL S.* Olso (FD01848867:15 FD01848867_02733) in pFPV25.1 |
| EBS 730-R HindIII | TTTTAAGCTTTCACTCCACTACCAGAATTTC | Clone *tssL S.* Olso (FD01848867:15 FD01848867_02733) in pFPV25.1 |
| EBS 879-F SacI | GAGCTCGCTGGATGTAATGCCCAAC | Clone SRLase1-SRLi1 in pDHL |
| EBS 880-R XmaI | CCCGGGTTATAATTTATCCATCCACGATCTCATG | Clone SRLase1-SRLi1 in pDHL |
| EBS 696-R SalI | AAAGTCGACCTATTTTTCAAATTGCGGATGGCTCCAAGCGCTCCCAGGAACGATAATTATCCCAG | Clone SRLase1-linker-Strep in pRSF-Duet MCS1 (FD01848867_02742), use with EBS 537-F |
| EBS 585-F NdeI | AATTATTCATATGAGAAATACAATAGATAATAGG | Clone *slri1* (FD01848867_02743) in MSC2 pRF-Duet |
| EBS 586-R KpnI | TAGGTACCTTATAATTTATCCATCCACGA | Clone slri1 (FD01848867_02743) in MSC2 pRF-Duet |
| EBS 313-F | GAACTGTGGGGAGTGCGCGATAAATGACAGACAGTGTGTAGGCTGGAGCT | Delete *tssL S.* Oslo |
| EBS 314-R | AGGTTGATTTTTTCATCATTCATTCACTCCACTACCATATGAATATCCTC | Delete *tssL S.* Oslo |
| EBS 324-F | CTATTACGACATCCGCCGC | Confirm *tssL S.* Oslo deletion |
| EBS 597-R | GCCACCGGGTAATAAACAAC | Confirm *tssL S.* Oslo deletion |
| EBS 598-F | CATGGATAGATCCGTTAGGCCTTGCTGGATGTAATGTGTAGGCTGGAGCT | Delete *srlase1/srli1* |
| EBS 599-R | AAATAAACCTCACAAAGATTTTATAATTTATCCATCATATGAATATCCTC | Delete *srlase1/srli1* |
| EBS 600-F | CAATTTGTACAGATTTTATGATCCGG | Confirm *srlase1/srli1* deletion |
| EBS 601-R | ATACCGGGGATATTCCGTTG | Confirm *srlase1/srli1* deletion |
| EBS 698-R FAM | FAM-ACCAGTGATGCGTCCACTCCG | Fluorescein tagged for primer extension assay |
| EBS 701-R FAM | FAM-ACCAGTGATGCGTCCACTCCGGTCCTCTCGTA | Fluorescein tagged ladder 32 nt |
| EBS 700-R FAM | FAM-AGGACCGGAGTGGACGCATCACTGGT | Fluorescein tagged ladder 26 nt |
| EBS 704-R FAM | FAM-AGGACCGGAGTGGACGCATCACTGG | Fluorescein tagged ladder 25 nt |
| EBS 705-R FAM | FAM-AGGACCGGAGTGGACGCATCACTG | Fluorescein tagged ladder 24 nt |
| EBS 706-R FAM | FAM-AGGACCGGAGTGGACGCATCACT | Fluorescein tagged ladder 23 nt |

[a]Restriction enzymes are underlined. Point mutations are in bold.

**Movie S1 (separate file).** Time-lapse microscopy of *E. coli* harboring pBRA SRLase1 growing in media supplemented with 0.5% D-glucose.

**Movie S2 (separate file).** Time-lapse microscopy of *E. coli* harboring pBRA SRLase1 growing in media supplemented with 0.2% L-arabinose.

**Dataset S1 (separate file).** SRLase1 homologs and their genomic context.
